# p63 and p73 regulate convergent and factor-specific transcriptional programs in cutaneous squamous cell carcinoma

**DOI:** 10.64898/2026.02.18.706563

**Authors:** Dario Antonini, Marco Ferniani, Claudia Russo, Marco Franciosi, Ludovica D’Auria, Sara Palumbo, Jieqiong Qu, Andrew P. South, Huiqing Zhou, Madhavi Kadakia, Gian Paolo Dotto, Caterina Missero

## Abstract

Aberrant transcriptional regulation is a defining feature of squamous cell carcinoma (SCC), yet how lineage transcription factors coordinate shared and factor-specific oncogenic programs remains poorly understood. Although *TP63* (p63) is frequently amplified in SCC, the contribution of its paralog *TP73* (p73) has remained unclear. Here we show that p73, together with p63, is upregulated in skin SCC and is required for tumorigenesis.

Mechanistically, p63 and p73 form heteromeric complexes and co-occupy distal enhancer elements, establishing a shared chromatin regulatory framework. Integration of chromatin and transcriptomic profiling reveals that this common enhancer landscape supports both convergent and divergent transcriptional outputs. Both factors cooperatively sustain core proliferation but also exert regulatory biases, with p63 preferentially reinforcing epithelial lineage circuits and p73 contributing to DNA replication and stress-associated pathways.

Among shared downstream targets, p63/p73 co-regulation of multiple epidermal growth factor receptor (EGFR) ligands establishes a feed-forward signaling module that amplifies mitogenic signaling. Amphiregulin emerges as a dominant functional mediator, and its depletion phenocopies key aspects of p63/p73 loss, including impaired proliferation and tumor formation.

Together, these findings support a model in which shared enhancer occupancy by p63 and p73 drives cooperative and factor-specific transcriptional programs, linking chromatin regulation to signaling and tumor maintenance.

## Introduction

Squamous cell carcinoma of the skin is primarily driven by chronic ultraviolet (UV) radiation exposure, which leads to the progressive accumulation of UV-signature mutations even in clinically normal sun-exposed skin (1–3). Therefore, the genetic architecture of skin SCC is complex, characterized by a high mutational burden, mutational heterogeneity, copy number alterations, and recurrent mutations in key tumor suppressor genes such as *NOTCH1*, *NOTCH2*, *TP53* (p53), *KMT2D* and *CDKN2A* (reviewed in (4)).

Network-based integrative analyses of TCGA tumors revealed that, despite mutational heterogeneity, a limited set of master regulators underlies cancer hallmarks (5), including all three p53 family members p53, p63, and p73. Both p63 and p73 genes give rise to multiple isoforms with distinct, sometimes opposing functions (6). Whereas TAp63 and TAp73 isoforms are generally low in abundance and function as tumor suppressors with pro-apoptotic activity, alternative truncated transcripts such as ΔNp63 and ΔNp73 are often overexpressed in cancer, where they promote cell proliferation and survival.

p63 is frequently amplified in SCCs, and ΔNp63 is commonly overexpressed through transcriptional activation, promoter hypomethylation, and/or increased protein stability, functioning as a central regulator of squamous lineage identity (7–13). Across SCC types, ΔNp63 promotes tumor initiation and maintenance by sustaining self-renewal, bypassing senescence, suppressing apoptosis, and reshaping differentiation, metabolic, and extracellular matrix programs (14–22). ΔNp63 overexpression typically coexists with p53 inactivation and, while it represses selected p53 target genes involved in cell-cycle arrest and apoptosis (23–26), it additionally executes a transcriptional program largely distinct from p53 (7,14,20,27–29). In mouse skin SCC, elevated p63 promotes oncogenic transformation, whereas genetic ablation of p63 leads to regression of established lesions (30–32).

Notably, transcriptomic analyses across the TCGA-defined pan-squamous transcriptional cluster indicate that p73 expression is elevated in SCCs arising from multiple anatomical sites (10). Consistently, proteomic analyses detect p73 expression in SCCs, with higher levels in common SCCs compared with rare SCC subtypes (33). Despite these omics data, the functional contribution of p73 to SCC transcriptional programs remains largely unexplored. In some HNSCC cell lines, TAp73 activates a pro-apoptotic transcriptional program that antagonizes ΔNp63α (17,34,35); however, the balance among p63 and p73 isoforms is critical in determining the transcriptional outcome. Importantly, p63 and p73 interact with each other and form heterooligomers due to their highly similar oligomerization domain, whereas wild-type p53 is unable to interact with either one (17,36,37). A heterotetramer composed of a p63 dimer and a p73 dimer is thermodynamically more stable than any other combination, including the two homotetramers (36,38). These p63-p73 heterotetramers have been detected *in vivo* in both human and mouse epithelial tissues, with elevated levels observed in human lung SCC (39).

EGFR-dependent signaling is a major contributor to squamous epithelial tumorigenesis, promoting proliferation, survival, and altered differentiation. Large-scale genomic analyses have identified EGFR amplification, and more rarely activating mutations, in approximately 15–20% of SCCs (40–42). In skin SCC, EGFR is frequently overexpressed and correlates with poor outcome in advanced and metastatic disease (43,44). EGFR activation in squamous tumors is mediated by multiple ligands expressed in squamous epithelia, including transforming growth factor-α (TGFA), betacellulin (BTC), heparin-binding EGF-like growth factor (HBEGF), amphiregulin (AREG), and epiregulin (EREG). Among these, AREG acts as a key autocrine factor for keratinocyte proliferation and is selectively upregulated in SCC, in parallel with increased EGFR phosphorylation (45–47). These findings support a model in which dysregulated expression of specific EGFR ligands cooperates with EGFR overexpression and downstream signaling to shape squamous tumor behavior.

Here, we investigate the functional interplay and specific contributions of p63 and p73 in skin SCC. By integrating genomic occupancy, transcriptomic profiling, and functional assays, we show that both transcription factors are required for skin SCC cell proliferation and tumorigenesis, while regulating overlapping as well as distinct sets of direct target genes. We identify multiple EGFR ligands as direct transcriptional targets of p63 and p73 and demonstrate that their expression is necessary to sustain SCC proliferation. Among these, AREG plays a central role in supporting tumor cell growth and tumorigenic potential.

## Results

### Distinct epidermal distribution and coordinated upregulation of p63 and p73 in SCC

Analysis of human tissue transcriptomes revealed that p73 is expressed in skin at levels higher than many other adult tissues (Figure S1A). We first compared p63 and p73 expression by semiquantitative immunofluorescence in a large cohort of normal skin, actinic keratosis (AK), and SCC. Consistent with previous reports ((48) and references therein), p63 displayed a strong nuclear signal in the basal and immediate suprabasal layers of the normal epidermis, with a gradual reduction in intensity in terminally differentiated layers (Figure 1A). By contrast, p73 expression was predominantly confined to keratinocytes directly anchored to the basement membrane, with detectable signal in most basal cells and only occasional suprabasal cells (Figure 1A). A similarly restricted expression pattern of p73 relative to p63 has been reported by the Human Pathology Atlas (49), not only in the epidermis but also in other stratified epithelia, including oral mucosa, esophagus, upper respiratory tract, and cervix. In these tissues, p73 expression is largely confined to basal cells, and to ciliated cells when present, consistent with its established role in multiciliogenesis (50,51). In line with these observations, quantitative proteomic analyses across a large cohort of SCCs originating from diverse tissues revealed that both p63 and p73 are expressed in typical SCCs, although p73 levels are, on average, lower than those of p63 (Figure S1B) (33). Notably, both proteins were significantly more highly expressed in conventional SCCs than in atypical SCCs.

**Figure 1.**
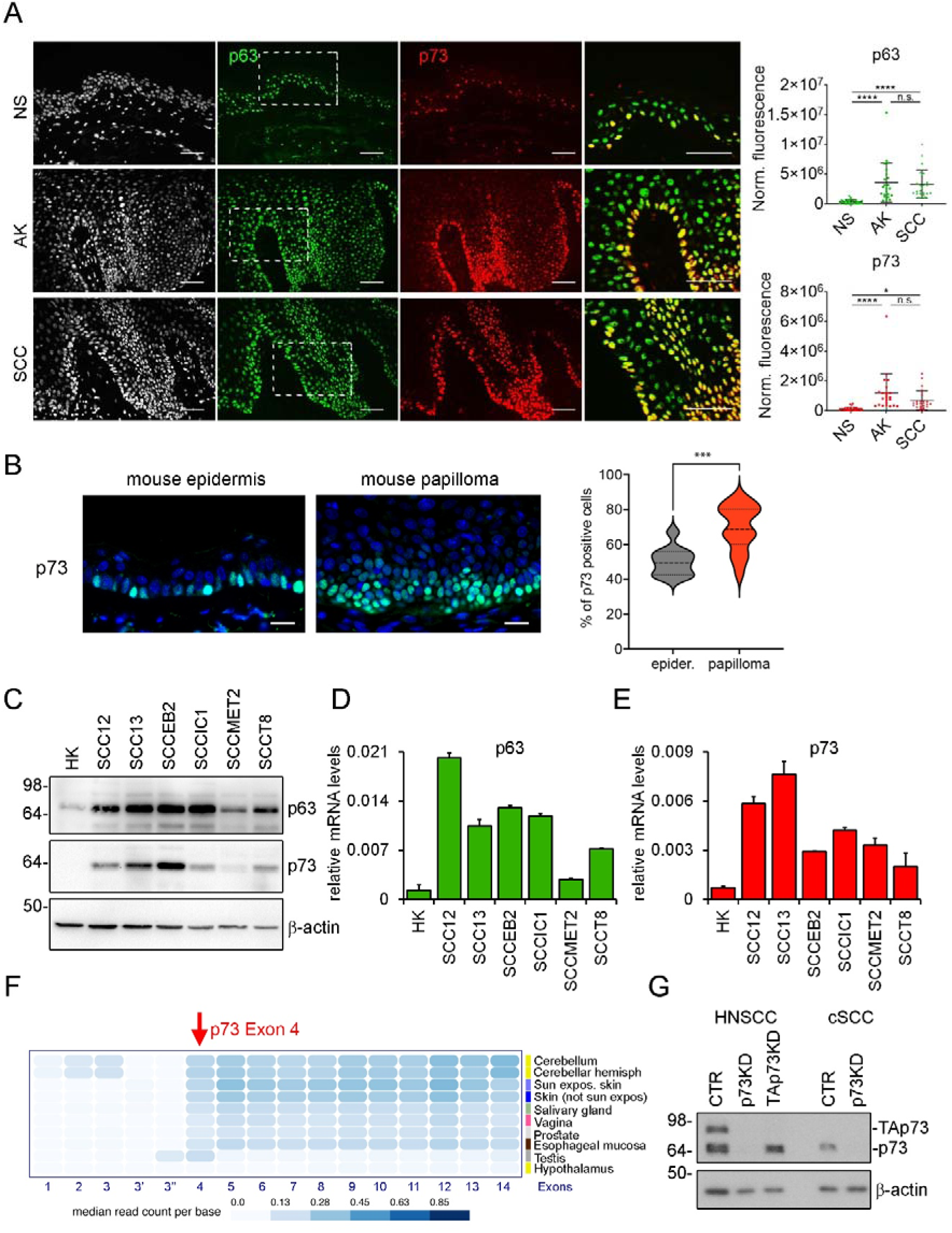
p63 and p73 expression in normal, precancerous, and malignant skin. (A) Immunofluorescence (IF) staining of normal skin (NS), actinic keratosis (AK), and SCC using antibodies against p63 (green) and p73 (red). Nuclei were counterstained with DAPI (white). Right panels show higher-magnification merged images of p63 and p73 co-staining. Scale bar, 50 μm. Quantification shows mean fluorescence intensity normalized to background (mean ± SD). Statistical analysis was performed by one-way ANOVA with Tukey’s multiple comparisons test. Sample numbers: NS n=28; AK n=23; SCC n=24. *P < 0.05; ****P < 0.0001. (B) IF staining of mouse adjacent epidermis and papillomas with anti-p73 (green); nuclei counterstained with DAPI (blue). Scale bar, 20 μm. Right: quantification of IF signal (pairwise t-test p = 0.037; n=4). (C) Immunoblot analysis of p63 and p73 protein expression in the indicated SCC-derived cell lines and in HK. (D-E) Quantification of total p63 (left) and p73 (right) mRNA expression in SCC-derived cell lines compared with HK. Data are presented as mean ± SD. (F) Heatmap showing median read counts per p73 exon across normal human tissues from the GTEx dataset. (G) p73 protein expression in SCC cells following transfection with pan-p73 or TAp73 siRNA; HNSCC cells (SCC011) expressing TAp73 were included as a control.

Both p63 and p73 proteins were markedly increased in the vast majority of AK and SCC compared with normal skin (Figure 1A). In these lesions, p73-positive nuclei were detected in nearly all basal keratinocytes, with additional clusters extending suprabasally. To further substantiate these findings, we analyzed a mouse model of chemically-induced skin carcinogenesis, in which papillomas similarly exhibited an increased number of p73-positive nuclei, predominantly within the basal compartment and extending into suprabasal cells, compared with the adjacent epidermis (Figure 1B).

Consistent with their *in vivo* expression patterns, p63 and p73 mRNA and protein levels were elevated in multiple human SCC-derived cell lines compared with primary human keratinocytes (HK), suggesting that their increased expression in tumors is not merely a consequence of expansion of the proliferative basal compartment (Figure 1C-1E). In both HK and in SCC, ΔNp63 was by far the predominant transcript, whereas TAp63 was barely detectable (Figure S1C).

Despite high levels of total p73 mRNA in SCC cells, previously described p73 transcripts, including TAp73, ΔNp73, Δex2, and Δex2/3, were not detected (Figure S1C). Instead, analysis of publicly available exon-specific datasets from the Genotype-Tissue Expression (GTEx) project and 5′-end RNA-seq (RAMPAGE) data from the ENCODE consortium indicated that skin predominantly expresses a 5′-truncated p73 transcript initiating at exon 4 (Figure 1F and S1D) (48,52). Consistent with this, selective depletion of TAp73 or ΔNp73 did not affect overall p73 expression in SCC cells, whereas pan-p73 depletion resulted in a marked reduction of p73 mRNA and protein levels (Figure 1G and S1E). As a control of TAp73-specific depletion, we used SCC011, a HNSCC cell line which exhibited abundant TAp73 expression as previously reported (17,24) (Figure 1G).

Together, these data establish that ΔNp63 is the dominant isoform in both normal and malignant keratinocytes, while p73 expression in skin is largely sustained by a truncated transcript initiating at exon 4, and that both transcription factors are consistently upregulated in premalignant and malignant skin squamous cancer.

### Both p63 and p73 are essential for cell proliferation and clonogenicity in SCC

To assess the biological significance of the elevated expression of p63 and p73 in SCC, we evaluated the effects of their depletion on cell proliferation in HK and SCC cells. Consistent with previous reports in HK and HNSCC (24,25), depletion of p63 or its ΔNp63 isoform markedly suppressed DNA synthesis, as assessed by BrdU incorporation, in four independent SCC cell lines as well as in HK (Figure 2A-C; Figure S2A). Notably, p73 depletion mirrored the effects of p63 loss, leading to a strong suppression of DNA synthesis in both HK and SCC cells (Figure 2A-C; Figure S2A). Flow cytometry of PI– and EdU/DRAQ5-stained cells showed that depletion of either p63 or p73 led to G1 accumulation in SCC (Figure 2D; Figure S2B), suggesting that both factors contribute to cell-cycle progression at the G1-S transition.

**Figure 2.**
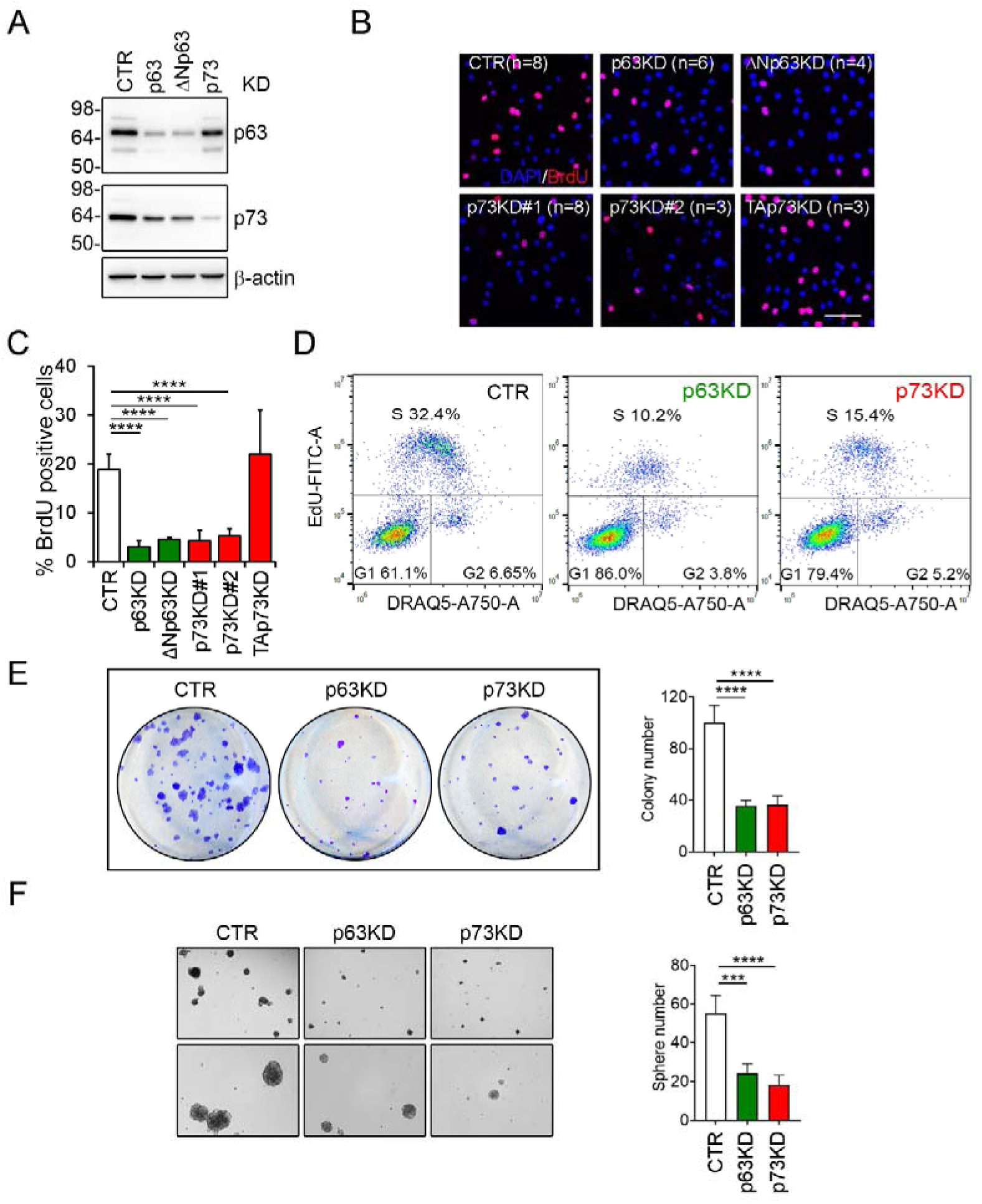
p63 and p73 are required for proliferation and clonogenic potential of SCC cells. (A) Immunoblot analysis of p63 and p73 protein levels in SCC13 cells following knockdown (KD) of p63, ΔNp63, or p73; β-actin was used as a loading control. (B) Representative IF images showing BrdU incorporation (red) in SCC13 cells 48 h after KD; nuclei were counterstained with DAPI (blue). Scale bar, 50 μm. (C) Quantification of BrdU-positive cells relative to total nuclei in SCC cells following the indicated KDs. Data are presented as mean ± SD. Statistical analysis was performed using two-tailed Student’s t-test (CTR, n=8; p63KD, n=6; ΔNp63KD, n=4; p73KD#1, n=8; p73KD#2, n=3; Tap73KD, n=2); ****P < 0.0001. (D) Cell cycle distribution of SCC cells following KD assessed by EdU incorporation and DNA content analysis by flow cytometry. (E) Clonogenic assay after KD. Left, representative crystal violet–stained colonies; right, quantification of colony number performed using ImageJ. Data are mean ± SD. One-way ANOVA with Tukey’s test; n=4; ****P < 0.0001. (F) Tumorsphere formation assay following KD. Left: representative images; right: sphere quantification using ImageJ. Data are mean ± SD. n = 4 *** P < 0.001, **** P < 0.0001.

We next assessed the long-term proliferative capacity of SCC cells following p63 or p73 depletion. Clonogenic assays showed a marked reduction in colony formation compared with controls (Figure 2E). Consistently, depletion of either factor impaired growth under anchorage-independent conditions, as indicated by reduced tumorsphere formation (Figure 2F). These findings indicate that both p63 and p73 are required to maintain long-term proliferative potential and/or to support the survival of tumor-initiating cell populations under non-adherent conditions.

To determine whether these growth defects were associated with increased cell death, we next examined apoptosis following p63 or p73 depletion. Annexin V staining and flow cytometric analysis failed to detect a significant increase in apoptotic cells upon depletion of either factor, in contrast to UV irradiation, which elicited a robust apoptotic response and served as a positive control (Figure S2C). These results suggest that the reduced clonogenic and tumorsphere-forming capacity observed upon p63 or p73 loss is unlikely to be explained by increased apoptotic cell death.

### Shared and factor-specific transcriptional programs regulated by p63 and p73 in SCC

To dissect how p63 and p73 shape the transcriptional landscape of SCC, we performed global gene expression profiling after depletion of each factor. Consistent with previous studies in HKs (53,54), p63 depletion markedly altered the transcriptome, with roughly equal numbers of genes being up– and downregulated (Figure 3A). Despite its lower abundance, p73 depletion affected a very similar number of genes with a comparable distribution (Figure 3A). The balanced pattern of up– and downregulated genes following depletion of either factor suggests that both p63 and p73 act as transcriptional activators as well as repressors in the SCC context. Notably, gene expression changes upon p63 or p73 depletion exhibited a robust positive correlation (r² = 0.731; Figure 3B), with nearly half of the affected genes co-regulated by both factors, underscoring their shared functions in SCC (Figure S3A).

**Figure 3.**
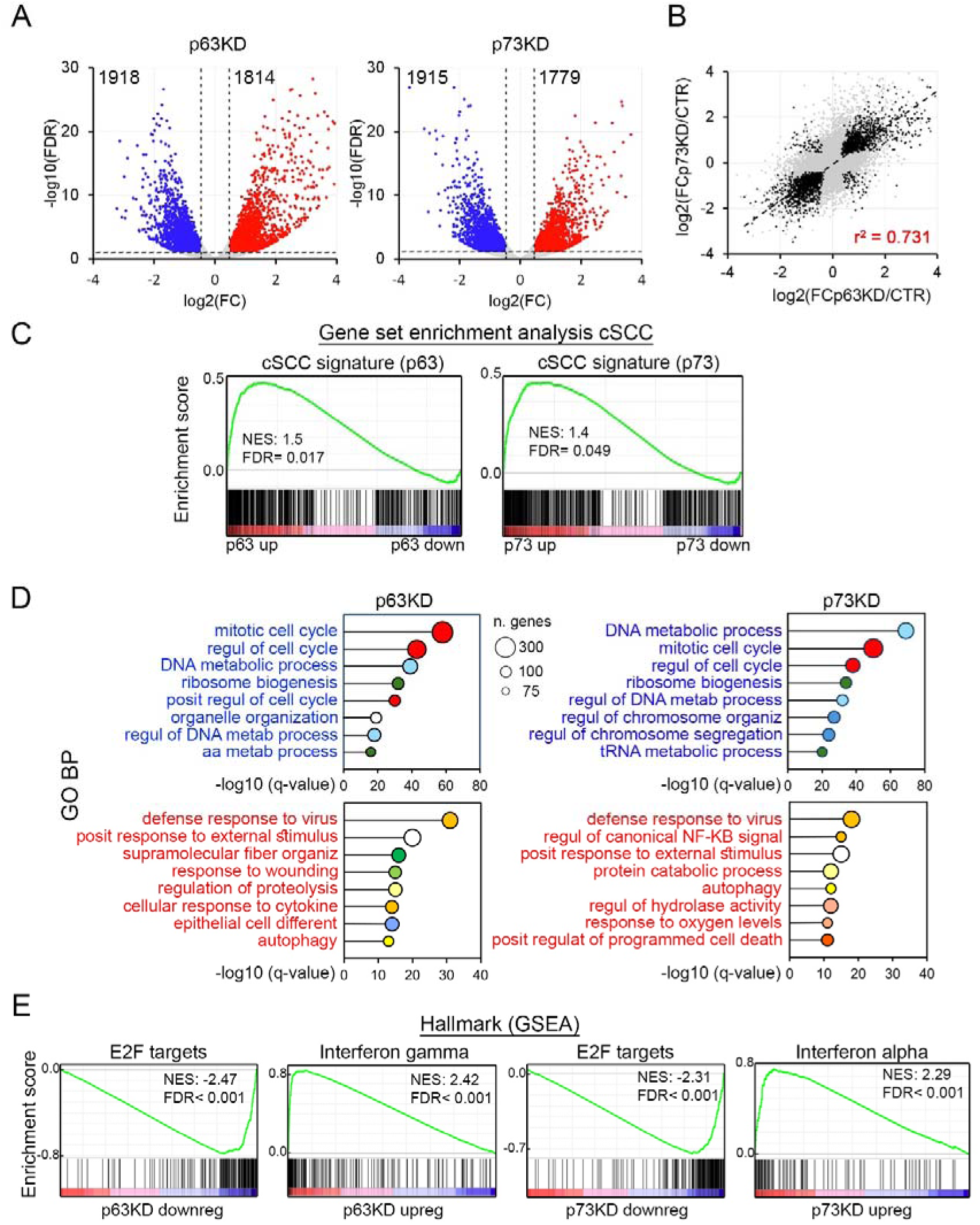
Integrated transcriptomic analysis of p63 and p73 function in SCC cells. (A) Volcano plots showing differences in RNA expression of p63KD (left) or p73KD (right) compared to control SCC cells. Thresholds: FDR < 0.05; log₂FC > 0.5; n=4. (B) Correlation analysis of gene expression changes following p63KD versus p73KD. Pearson correlation coefficient and linear regression line are indicated. (C) GSEA showing enrichment of an SCC transcriptional signature (PRJNA844527) (56) in p63– and p73– KD SCC cells. NES, normalized enrichment score. (D) Lollipop plots representing the enrichment of biological processes (GO BP) of upregulated (text in red) and downregulated (text in blue) genes in p63– (left panel) or p73– (right panel) depleted SCC cells compared to controls. Circle size indicates gene count. Circle colors denote functionally related GO terms. X-axis shows −log₁₀FDR. (E) GSEA of selected hallmark gene sets enriched among genes differentially expressed after p63 or p73 depletion. NES, normalized enrichment score

Gene Set Enrichment Analysis (GSEA) (55) revealed a strong enrichment of p63– and p73-regulated genes in the SCC transcriptional signature derived from a large patient-based dataset comparing SCC with normal skin (56) (Figure 3C). This supports a role for both p63 and p73 in enforcing SCC-associated transcriptional programs.

Genes downregulated upon depletion of either p63 or p73 were significantly enriched for biological processes related to cell cycle regulation and DNA metabolism (Figure 3D), in line with the proliferation defects observed in SCC cells lacking these factors. Hallmark GSEA further identified prominent enrichment of E2F and MYC target gene sets, two central regulators of proliferative transcriptional programs and cell cycle progression (Figure 3E and S3B). Conversely, genes upregulated upon depletion of p63 or p73 were enriched for pathways related to interferon signaling, inflammatory responses, and apoptosis (Figure 3D-E, S3B). Together, these data indicate that loss of either factor activates a pro-inflammatory transcriptional program, accompanied by induction of cell death–associated gene signatures in the absence of overt apoptosis (Figure 3D-E and Figure S3B).

Despite the extensive overlap, each factor also controlled distinct gene sets. Genes preferentially inhibited by p63 deletion included well-established p63 targets (Figure S3C). By contrast, p73 depletion predominantly inhibited genes involved in DNA replication and replication stress (Figure S3C). These findings suggest that while p63 primarily reinforces epithelial integrity and lineage fidelity, p73 contributes more prominently to DNA replication and repair.

Consistent with the well-characterized function of p63 in the epidermis, genes selectively induced by p63 depletion were enriched for late epidermal differentiation (Figure S3C). In contrast, genes upregulated upon p73 depletion were associated with apoptotic signals in response to DNA damage (Figure S3C).

Together, these findings indicate that p63 and p73 co-regulate core transcriptional programs, including cell cycle control and inflammatory responses, while also governing partially distinct gene networks.

### Extensive co-occupancy of p63 and p73 at enhancer regions in SCC

Given the extensive overlap between genes affected by p63 and p73 depletion (Figure S3A), we next examined whether the two proteins physically interact in SCC cells. Structural studies have shown that the tetramerization domains of p63 and p73 can assemble into stable heterotetramers (36,38,57), and endogenous p63-p73 complexes have been described previously in keratinocytes and in HNSCC cells (17,36). Proximity ligation assays demonstrated close nuclear association of p63 and p73 in SCC cells (Figure 4A). Co-immunoprecipitation with antibodies against either factor further demonstrated a robust endogenous interaction between p63 and p73 (Figure 4B). These findings support the coexistence of p63/p73 heteromeric complexes together with p63 homomeric complexes.

**Figure 4.**
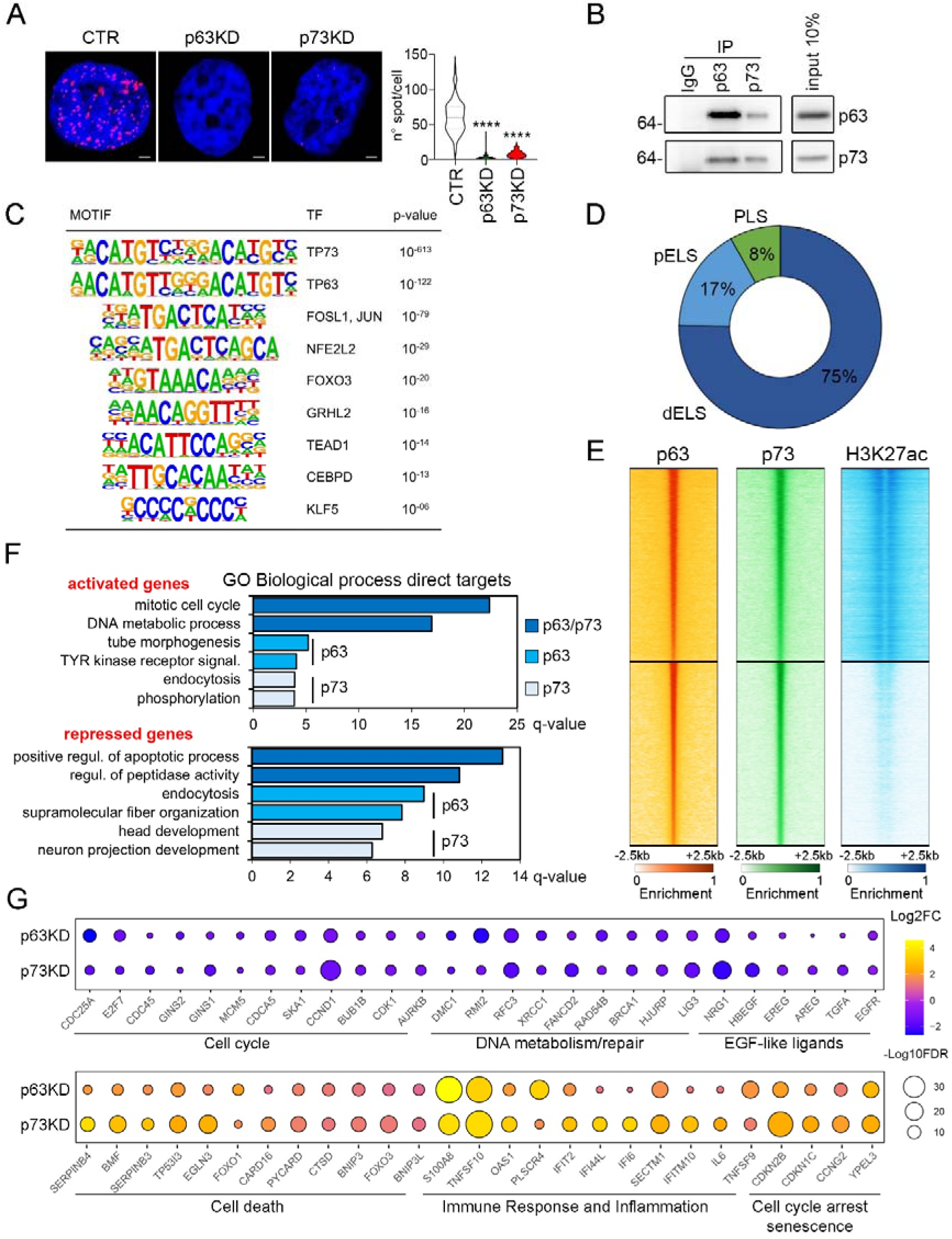
p63 and p73 form endogenous complexes and co-occupy distal enhancers that regulate epithelial and SCC-specific gene programs. (A) PLA showing nuclear colocalization of endogenous p63 and p73 in SCC cells. Representative images of control, p63KD, and p73KD cells, and quantification of PLA signal per nucleus (right). Scale bar: 2 µm. Statistical significance was performed using one-way ANOVA and Tukey’s multiple comparisons test. Number nuclei: CTR, n=79; p63KD n=122; p73KD n=153. **** P < 0.0001. (B) Co-immunoprecipitation of endogenous p63 and p73 in SCC cells using the indicated antibodies followed by immunoblot detection. (C) Top enriched DNA-binding motifs within p63– and p73-bound regions identified by ChIP-seq. A centrally positioned p63/p73 consensus motif is the most enriched, followed by the consensus for the indicated transcription factors. (D) Genomic annotation of p63/p73 co-bound regions showing enrichment at distal enhancer-like signatures (dELS) relative to promoter-like signatures (pELS) and other regulatory elements. (E) Heatmaps of p63, p73, and H3K27ac ChIP-seq signal centered on consensus p63/p73 motifs, indicating overlap with active enhancer marks. (F) GO Biological Process enrichment analysis of genes directly associated with p63/p73 co-bound genomic regions and transcriptionally regulated by both factors. (G) Balloon plot of representative p63– and p73-regulated genes grouped by biological function. Bubble size represents log₂ fold change; color intensity indicates −log₁₀ FDR.

To determine whether p63 and p73 occupy overlapping genomic regions, we performed chromatin immunoprecipitation followed by sequencing (ChIP-seq). The two factors displayed extensive co-occupancy, with p73 binding largely coinciding with p63 peaks but at markedly lower levels (on average one-third of the p63 signal) (Figure S4A), consistent with its lower abundance. As expected, a centrally positioned p63/p73 consensus motif was the most enriched element within both p63– and p73-bound regions (Figure 4C). Motifs for AP-1 family members were also strongly enriched, consistent with our previous findings and independent studies (7,54,58–61). In addition, motifs characteristic of p63-regulated enhancers, including GRHL2, TEAD1, KLF4/5, and CEBPD, were significantly enriched (7,60–63). Notably, motifs for FOXO3 and NFE2L2 were among the most enriched, despite not being previously detected in p63 ChIP-seq datasets. This observation is consistent with previous evidence of molecular interactions between p63 and FOXO proteins (64) as well as between p63 and NFE2L2 (65). Together, these findings suggest that such factors may contribute more broadly to enhancer programs co-regulated by p63 and p73.

The majority of p63/p73 binding sites localized to distal genomic regions and to non-first-intron intronic genomic regions, consistent with enhancer elements (Figure S4B) and overlapped extensively with ENCODE-defined candidate cis-regulatory elements (cCREs) (66) (Figure 4D). Many target genes were associated with more than one such region, with 30% containing three or more p63/p73 binding sites, underscoring the dense regulatory architecture around these loci. Genes linked to p63/p73 binding regions were significantly enriched for biological processes central to keratinocyte identity, including epidermis development and epithelial differentiation (Figure S4C), and were also strongly associated with cancer-related pathways and MAPK signaling components (Figure S4D).

To determine which p63/p73 binding regions correspond to active enhancers or promoters in SCC, we intersected them with H3K27Ac-enriched regions previously mapped in SCC (67). A substantial fraction of p63/p73 binding sites overlapped with H3K27Ac, consistent with their localization at active regulatory elements (Figure 4E and S4E). Genes linked to H3K27Ac-positive regions (6,342 genes) were strongly enriched for p63 binding in human keratinocytes (HK), based on transcription factor enrichment analysis using the ChEA 2022 database (adjusted p = 2.05 × 10^-260^). In contrast, genes associated with H3K27Ac-negative p63/p73-bound regions (2,123 genes) showed markedly weaker enrichment for p63 binding in HK (adjusted p = 2.82 × 10^-10^). Notably, genes linked to H3K27Ac-flanked p63/p73 sites were preferentially associated with keratinocyte identity, whereas those linked to p63/p73 binding sites lacking H3K27Ac were enriched for neuronal signatures (Figure S4F). Together, these observations suggest that p63 and p73 contribute to the suppression of neuronal gene programs in SCC, consistent with prior observations in p63-depleted mouse keratinocytes and during zebrafish ectoderm specification (61,68,69).

The vast majority of p63/p73 binding regions identified in SCC overlapped with sites previously mapped in neonatal HK (72%) and adult HK (81%) (53,54,70), and more than half were also shared with oral SCC (57%) (32). A direct comparison revealed a distinct subset of p63/p73 binding sites present in skin-derived keratinocytes (newborn HK, HK, SCC) and absent in OSCC, indicating epithelial lineage-specific differences in enhancer usage. Genes associated with these skin-restricted sites were strongly enriched for processes such as epidermis development and neuron projection, in line with the well-established role of p63 as a key regulator of ectodermal specification.

A notable example is the pair of evolutionarily conserved enhancer regions C38 and C40, located within a large intron of the p63 gene. These elements are bound by p63 and control tissue-and layer-specific p63 expression during mouse skin development. C40 exhibits strong epithelial-specific activity and functions as a broad regulator of p63 expression (60,71,72), whereas C38 lacks intrinsic enhancer activity but cooperates with C40 to modulate p63 regulation during keratinocyte terminal differentiation (72). In line with these properties, p63 binding to C40 was detected across all examined cell types, while C38 binding was restricted to epidermal keratinocytes and SCC and absent in OSCC, consistent with its skin-specific regulatory role (Figure S4G, left panel).

Interestingly, a small subset of p63/p73 binding sites was unique to SCCs (SCC and OSCC) and absent in keratinocytes. These SCC-specific regions were associated with genes enriched for leukocyte activation and cytokine production, consistent with the inflammatory signature observed in the global expression analysis (Figure 3D-E and S3B). In addition to these immune-related loci, this SCC-restricted set also included genes with well-established roles in squamous tumor biology. A notable example is ACTL6A (Figure S4G, right panel), a core subunit of the SWI/SNF chromatin remodeling complex frequently amplified in SCCs. p63/p73 binding to ACTL6A was detected only in SCCs and not in keratinocytes, suggesting that SCC-specific enhancer engagement may contribute to its elevated expression and its previously demonstrated requirement for SCC proliferation (7). These findings indicate that p63 and p73 engage enhancer landscapes in a cell type-dependent manner, thereby shaping tissue-specific gene regulatory programs. Although many p63 target genes are flanked by multiple p63/p73 binding regions, the coordinated use of these sites likely provides an additional layer of regulatory flexibility, enabling precise modulation of transcriptional outputs across epithelial contexts.

### Direct transcriptional targets of p63 and p73 in SCC

To identify directly regulated genes of p63 and p73 in SCC, we integrated RNA-seq data with ChIP-seq binding profiles. This analysis uncovered approximately 2,200 genes directly regulated by either factor (FDR < 0.05, log₂ Fold Change (FC) > 0.5). These genes were nearly evenly divided between those downregulated and upregulated upon depletion, indicating that both p63 and p73 function as transcriptional activators and repressors in SCC. Notably, 45% of the directly regulated genes were affected in the same direction by both factors, highlighting a substantial degree of functional overlap between p63 and p73 in shaping the transcriptional landscape of SCC cells.

Consistent with the cell cycle arrest observed in SCC cells following p63 or p73 depletion, a substantial fraction of the directly downregulated targets comprised genes involved in cell-cycle progression, DNA metabolism and repair, and the EGFR signaling (see below), many of which had not been previously recognized as direct p63 or p73 targets (Figure 4F-G).

In contrast, genes directly upregulated upon p63/p73 depletion were enriched for pathways associated with cell death, cellular stress responses, inflammation, and cell-cycle arrest (Figure 4F-G). Among these were FOXO3 and, to a lesser extent, FOXO1, key stress-responsive transcription factors activated under conditions such as reduced growth factor signaling (73). We previously showed that impaired IGF1 signaling promotes FOXO nuclear accumulation and suppresses p63 activity in mice (64), suggesting a negative feedback loop between p63/p73 and FOXO factors. Consistent with this model, prominent p63– and p73-binding regions were detected within the first intron of both FOXO1 and FOXO3 in multiple epithelial cell types, indicating a conserved regulatory interaction (Figure S5A).

To identify pathways preferentially controlled by each factor, we examined direct targets predominantly affected by p63 or p73 depletion. Genes downregulated after p63 depletion (i.e., activated by p63) included several well-known p63 targets involved in epithelial morphogenesis and function, NOTCH signaling, tyrosine kinase receptors (e.g. COL7A1, FGFR2, NOTCH1, JAG1), as well as genes linked to energy metabolism and oxidative stress (Figure S5B). Genes predominantly downregulated following p73 depletion were enriched for endocytosis, DNA metabolic processes, replication and repair (e.g., PARP1, POLE4, POLQ), and also included several genes involved in cilium assembly, consistent with the established role of p73 in ciliogenesis (50,51,74) (Figure S5B).

Genes upregulated after p63 depletion (i.e., repressed by p63) were enriched for keratinocyte differentiation genes (e.g. KLK5, TGM1, SCEL, MAFB), cytoskeletal organization, and extracellular matrix remodeling (COL5A1, VIM, various MMPs) (Figure S5B), consistent with p63’s role in maintaining the basal keratinocyte program. p63 also repressed several immune and inflammatory genes, including components of the STING pathway (IF16, TREX1, UNC93B1). In contrast, genes upregulated after p73 depletion were associated with nervous system development, and canonical p53-family targets (e.g. TNFRSF10D, SESN1, CLCA2) (Figure S5B).

Together, these findings show that p63 and p73 jointly sustain core proliferative and stress-response programs in SCC, while their factor-specific activities extend this shared network to fine-tune epithelial identity and tumor behavior.

### Cooperative transcriptional control of EGFR ligands by p63 and p73 sustains EGFR signaling in SCC

Among the transcriptional programs jointly regulated by p63 and p73, multiple components of the EGFR pathway were consistently downregulated upon depletion of either factor, prompting us to examine whether cooperative regulation of EGFR ligands represents a key mechanism by which p63 and p73 sustain SCC proliferation. EGFR signaling provides a well-established growth advantage in cancer by activating canonical downstream pathways (75). Consistent with this, integrated multi-omic analyses revealed that p63 and p73 regulate multiple components of the EGFR and MAP kinase pathways. Notably, several EGFR ligands (AREG, EREG, HBEGF, and TGFA) were downregulated at the mRNA level upon p63 or p73 depletion in several SCC cell lines, supporting a positive role for both factors in EGFR ligand expression (Figures 5A and S6A). Among these, AREG was the most abundantly expressed, consistent with its autocrine function in keratinocyte proliferation (46,47,76). Reduced AREG secretion upon p63 depletion was confirmed by ELISA (Figure 5B). In parallel, depletion of p63 or p73 was associated with reduced levels of phosphorylated EGFR and, to a lesser extent, total EGFR protein (Figure S6B).

**Figure 5:**
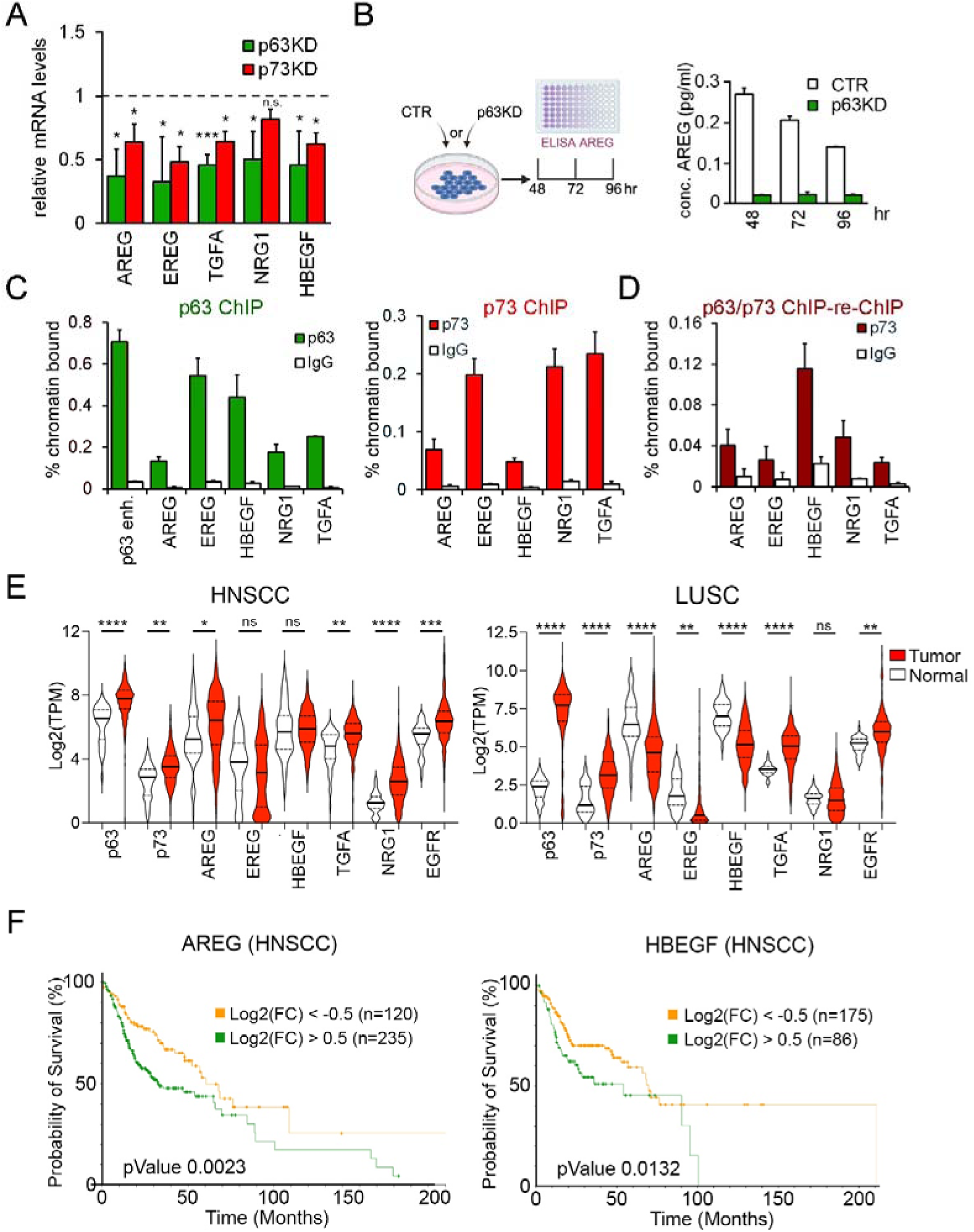
p63 and p73 control cell proliferation in SCC cells through EGF signaling pathway. (A) RT-qPCR analysis of EGFR ligand mRNA levels in SCC cells after p63 or p73 KD. Data are mean ± SD relative to controls. Two-tailed Student’s t-test; n=6; *P < 0.05; ***P < 0.001. (B) Schematic of ELISA workflow (left) and quantification of secreted AREG protein in conditioned media from control or p63KD SCC cells (right). (C) ChIP-qPCR showing p63 or p73 occupancy at EGFR ligand regulatory regions in SCC13 cells. (D) Sequential ChIP (re-ChIP) demonstrating co-occupancy of p63 and p73 at the indicated loci. (E) Violin plots of p63, p73, EGFR ligands, and EGFR expression in tumor versus normal samples from HNSCC and LUSC cohorts (TCGA). Values represent log₂(TPM). One-way ANOVA with Tukey’s test. Sample numbers: HNSCC normal n=44, tumor n=502; LUSC normal n=49, tumor n=501. (F) Kaplan–Meier survival analysis of HNSCC patients (TCGA) stratified by AREG or HBEGF expression. p values are indicated.

Analysis of the genomic loci of EGFR ligands identified multiple regulatory regions co-occupied by p63 and p73 annotated as active enhancers (Figure S6C), consistent with direct transcriptional regulation. Binding of both factors to the most enriched regulatory element associated with each ligand gene was validated by ChIP-qPCR (Figures 5C). Sequential ChIP (Re-ChIP) assays further demonstrated simultaneous occupancy of p63 and p73 at the same chromatin regions, supporting cooperative regulation of EGFR ligand genes (Figure 5D).

Consistent with these findings, analysis of TCGA head and neck and lung SCC datasets revealed coordinated upregulation of p63, p73, EGFR, and multiple EGFR ligands, including AREG, TGFA, and NRG1, in tumors compared with normal tissues (Figure 5E). Notably, high AREG expression was associated with significantly poorer overall survival (Figure 5F), consistent with a pro-tumorigenic role for EGFR ligand signaling in SCC.

To assess the functional relevance of EGFR signaling, EGFR or AREG were individually depleted by siRNA (Figure S6D-E). Depletion of either gene significantly impaired cell proliferation, as measured by BrdU incorporation, and markedly reduced clonogenic and anchorage-independent growth (Figure 6A-C). Conversely, treatment with recombinant AREG or HBEGF restored ERK and AKT phosphorylation in both p63– and p73-depleted cells (Figure 6D). Functionally, ligand supplementation robustly rescued BrdU incorporation in p63-depleted cells, and also improved proliferation in p73-depleted cells, with a significant effect observed in AREG-treated cultures (Figure 6E). These data indicate that EGFR ligand signaling is sufficient to restore proliferation downstream of p63 loss and can also support proliferative recovery in p73-depleted cells, albeit to a more limited extent. Finally, to test the requirement of the p63/p73-EGFR ligand axis for SCC tumorigenicity *in vivo*, control and depleted SCC cells were injected subepidermally to preserve the native skin microenvironment (Figure 6F, S6F). Control cells formed rapidly expanding tumors, whereas depletion of p63 or p73 profoundly impaired tumor formation. Notably, AREG depletion similarly abolished *in vivo* tumor growth (Figure 6F, S6F). Collectively, these findings demonstrate that p63 and p73 cooperate to sustain EGFR ligand expression, and in particular AREG, thereby maintaining EGFR signaling required for SCC proliferation and tumorigenesis.

**Figure 6:**
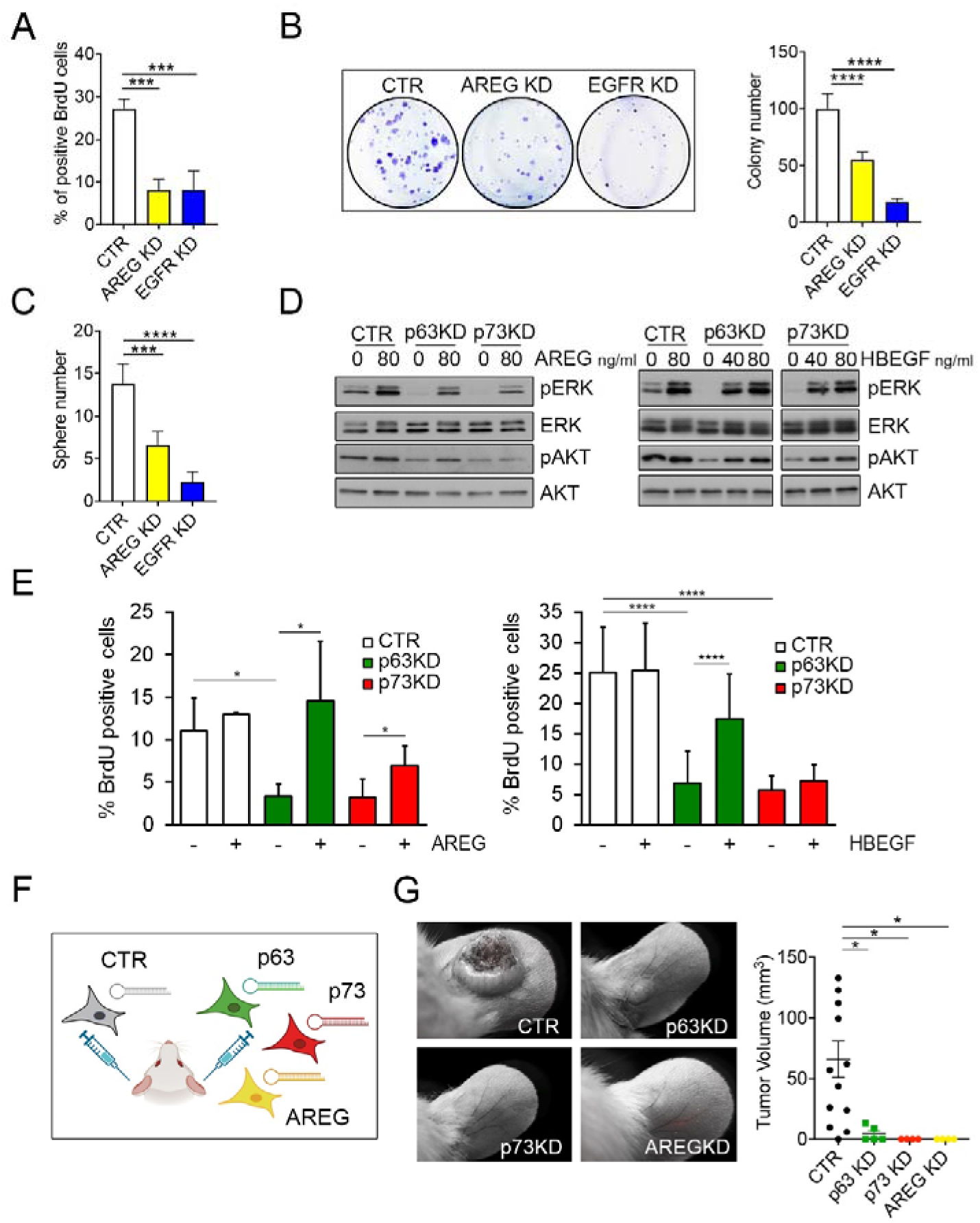
Functional and clinical relevance of p63/p73-EGFR ligand axis in squamous cell carcinoma. (A) BrdU incorporation in SCC cells transfected with the indicated siRNAs. Data are mean ± SD. One-way ANOVA with Tukey’s test; n=3; ***P < 0.001. (B) Clonogenic growth of SCC13 cells after siRNA transfection. Representative colonies and quantification are shown. Mean ± SD; one-way ANOVA with Tukey’s test; n=4; ****P < 0.0001. (C) Tumorsphere formation following siRNA transfection. Sphere number quantified with ImageJ. Mean ± SD; one-way ANOVA with Tukey’s test; n=4; ***P < 0.001; ****P < 0.0001. (D) Immunoblot analysis of signaling pathway activation following AREG or HBEGF treatment in SCC cells transfected with the indicated siRNAs. (E) BrdU incorporation after AREG or HBEGF stimulation. Mean ± SD. Two-tailed Student’s t-test; left panel n=3, right panel n=11. *P ≤ 0.01; ***P ≤ 0.0001. (F) Schematic of the in vivo tumorigenicity assay. SCC cells expressing control or shRNAs targeting p63, p73, or AREG were injected subepidermally into immunodeficient mice. (G) Representative tumors and endpoint tumor volume quantification. Each dot represents one tumor; bars indicate mean ± SD. One-way ANOVA with Tukey’s test. CTR n=13; p63KD n=5; p73KD n=4; AREGKD n=4.

## Discussion

In this study, we show that the two p53 family members p63 and p73 are both highly expressed in SCC and are required for maintaining its proliferative potential and tumor-initiating capacity. Although neither factor is frequently mutated in cancer, both have been classified as cancer master regulators, integrating the heterogeneous effects of mutations and aberrant upstream signaling into a transcriptional program that establishes and sustains the malignant cell state (5).

The basal layer of many stratified epithelia expresses both p63 and p73, but their shared and distinct functions in these tissues, particularly in squamous cancers, remain poorly defined. In particular, the contribution of p73 to SCC biology is essentially unexplored. Here, we show that the two factors bind the same genomic regions in SCC, without detectable separation of their cistromes. They operate together to control an extensive set of genes involved in many aspects of cell functions promoting cell-cycle progression, DNA metabolism, and suppressing inflammatory pathways. This coordinated activity reflects their ability to recognize identical consensus motifs and to co-occupy distal enhancers and more rarely proximal promoters, thereby driving a common oncogenic transcriptional program.

The functional divergence between the two factors becomes apparent upon depletion. Although ChIP-seq reveals widespread co-binding, depletion experiments expose factor-selective dependencies. A major biological program under strong positive control by both p63 and p73 is cell proliferation. Both promote this process by directly regulating genes involved in cell-cycle progression and DNA metabolism, and by enhancing upstream mitogenic signaling, including the EGFR/MAPK pathway. Nevertheless, we observed clear pathway biases. p63 preferentially sustains receptor tyrosine kinases and genes implicated in epidermal development, differentiation, and NOTCH signaling, consistent with its established role as a lineage determinant. In contrast, p73 more strongly regulates genes involved in centrosome organization, ciliogenesis, and selected enzymatic components of DNA replication, repair, and chromatin regulation. These distinctions indicate that differences within a largely shared cistrome translate into selective transcriptional outputs that are revealed only when each factor is individually perturbed.

Mechanistically, the transcriptional outcome at shared regulatory regions is likely governed by the stoichiometry and isoform composition of p63 and p73 within individual cells. Biochemical studies have shown that the p63 and p73 oligomerization domains can assemble into several heterotetrameric configurations, including dimers of homodimers, which are thermodynamically more stable than the corresponding homotetramers, as well as other, less stable mixed arrangements (36,38,57). Although dimers of homodimers represent the most stable configuration, tetramer composition *in vivo* will depend on the relative abundance of each protein. In SCC, p63 is substantially more abundant than p73, suggesting that heterotetramers containing limited amounts of p73 may predominate at shared binding sites. In a physiological context where p73 levels fluctuate across basal subpopulations, even modest changes in p63/p73 ratio could alter DNA-binding stability, cofactor recruitment, or chromatin remodeling, thereby shifting transcriptional output in a graded manner.

A further level of complexity arises from isoform diversity. In SCC, as in primary keratinocytes, the predominant p63 isoforms are ΔNp63α and ΔNp63β. By contrast, the most abundant p73 isoform lacks both the TA and ΔN N-terminal domains, and therefore the canonical transactivation domains. Structural differences at the N-terminus may contribute to gene-selective transcriptional effects, potentially by modulating interaction surfaces for co-activators or chromatin regulators. Importantly, our integrated multi-omics analysis argues against a model in which p73 functions solely as a dominant-negative or repressor in SCC, indicating instead a context-dependent and cooperative contribution to transcriptional activation. Interestingly, while our study was ongoing, Weinberg and colleagues independently examined p63/p73 function in metastatic breast cancer cells (77,78). In that context, ΔNp63 is expressed at relatively low levels, consistent with cells transitioning away from a fully epithelial identity, and two major p73 isoforms, including TAp73, are present. Consequently, the p63/p73 isoform balance differs markedly from that in SCC, where TAp73 is undetectable and high levels of ΔNp63 are essential to sustain the squamous transcriptional program. Despite these differences in isoform composition and lineage state, both systems converge on a similar transcriptional outcome: joint activation of EGFR ligands. This convergence suggests that distinct isoform combinations of p63 and p73 can cooperate to engage overlapping oncogenic signaling nodes, and that the longstanding model in which TA– and ΔN-isoforms simply antagonize each other is likely an oversimplification, as suggested also in other cellular contexts (reviewed in (79)).

Among the shared downstream effectors that translate p63/p73 chromatin co-occupancy into oncogenic signaling, the EGFR pathway represents a major downstream effector pathway. Here we show that several EGF-family ligands—including AREG, EREG, HBEGF, and TGFA—are directly regulated by both p63 and p73.Consistent with the selective enrichment of EGFR ligands in SCC, EGFR phosphorylation is higher in bulk SCC compared with BCC or normal skin, and transcripts for AREG, HBEGF, and TGFA are specifically upregulated in SCC tumor cells but not in BCC (45). In our study, among the EGFR ligands induced by p63/p73, AREG emerges as the dominant paracrine mediator linking their activity to EGFR pathway activation in SCC.

More broadly, a comprehensive mapping of oncogene dependencies has shown that EGFR dependence in lung and other solid tumors is driven not only by EGFR amplification or high receptor expression but also by elevated AREG levels, which enhance EGFR addiction and downstream signaling (80). Elevated AREG expression is associated with worse prognosis in HNSCC; however, ligand-high tumors often display increased sensitivity to anti-EGFR therapy, and AREG and EREG expression can predict better responses to combinations of cetuximab and chemotherapy in recurrent or metastatic disease (81,82). Similar ligand-response associations have been reported in gastroesophageal and advanced colorectal cancers (83–85), underscoring the functional relevance of EGFR ligands across diverse epithelial malignancies.

Advanced SCC is frequently not amenable to curative surgery or radiotherapy, and conventional chemotherapy provides limited clinical benefit. Immune checkpoint inhibitors, such as the anti-PD-1 antibody cemiplimab, represent the standard systemic therapy for advanced disease due to higher and more durable response rates (86–88); however, a substantial fraction of patients fail to respond or eventually relapse. EGFR-directed therapy with monoclonal antibodies such as cetuximab therefore remains clinically relevant (reviewed in (89)), particularly in PD-1-refractory settings and in combination regimens, despite variable efficacy and increased toxicity (90).

Importantly, cetuximab acts primarily by blocking ligand-dependent EGFR activation rather than constitutive receptor signaling. In this context, our data provide a mechanistic framework to explain the heterogeneous clinical responses to EGFR inhibition in SCC. By demonstrating that p63 and p73 cooperatively sustain expression of specific EGFR ligands, we identify ligand-driven EGFR signaling as a biologically defined vulnerability in a subset of SCCs. These findings suggest that assessing EGFR-ligand expression, rather than EGFR levels alone, may represent a clinically informative strategy to predict sensitivity to anti-EGFR therapies and provides a rationale for developing novel ligand blockers to rationalize their use in advanced or refractory disease.

These observations may extend beyond SCC to other tumor types that express p63 and p73, including pancreatic cancer. In this context, p63 drives the adenocarcinoma-to-squamous transition, a switch associated with markedly worse outcomes (91,92), while p73 is expressed in basal-like pancreatic tumors and activates transcriptional programs linked to basal identity and ciliogenesis (93). Notably, AREG is strongly upregulated in pancreatic tumors, cancer cell lines, and cancer-associated fibroblasts, where it promotes metastatic progression (94–97). Together, these data support the testable hypothesis that p63 and p73 may contribute to AREG regulation in pancreatic cancer, with functional consequences for tumor progression.

## Methods

### Cell culture, transfections and infections

Skin-derived cells were maintained at 37°C in 5% CO2 in Keratinocytes-SFM medium supplemented with Bovine Pituitary Extract, recombinant human Epidermal Growth Factor (Thermo Fisher, 17005042) and penicillin/streptomycin solution. HNSCC were maintained at 37°C in 5% CO2 in DMEM with 10% FBS and penicillin/streptomycin solution. For depletion experiments, cells were transfected with specific silencing RNA or control siRNA (Thermo Fisher, 12935300) at a final concentration of 100nM using Lipofectamine 2000 (Thermo Fisher, 11668019), following the manufacturer’s protocol.

For lentiviral preparation, HEK293T cells were cultured at 37°C in 5% CO2 in DMEM, supplemented with L-glutamine, 10% FBS and penicillin/streptomycin solution and co-transfected with lentiviral packaging plasmids and pLKO.1 puro (Gift from Bob Weinberg; Addgene plasmid # 8453), containing the short hairpin RNA of interest using Lipofectamine 2000. Lentiviral particles were collected after 48-72 hr. SCC cells were infected for 2 hr. Two days after infection, cells were selected with 1 μg/ml of puromycin (Sigma-Aldrich, P8833) for 3 days.

### Clonogenic and sphere assays

For clonogenic assays, cells were plated 24 hr after transfection at a density of 500 cells per 60-mm dish and cultured in supplemented Keratinocyte-SFM for 10 days. The medium was refreshed every 2 days. After 10 days, colonies were fixed in 4% formaldehyde in PBS and stained with 1% crystal violet in 80% methanol. Colonies were counted and analyzed using ImageJ.

For sphere assays, cells were trypsinized, centrifuged, and resuspended in spheroid medium (DMEM/F12) supplemented with 2% B27 serum-free supplement (Thermo Fisher, 17504-044), 0.4% BSA, 20 ng/ml EGF (Sigma-Aldrich, E4269), µg/ml insulin, (Sigma-Aldrich, 19278) as previously described (13). A total of 40,000 cells were plated in each ultra-low attachment 35 mm-well and allowed to grow for 10 days.

### Mouse studies

Mouse skin tumors were generated in three-month-old male C57BL/6 mice using a long-term chemical carcinogenesis protocol with DMBA (9,10-dimethyl-1,2-benzanthracene) (Selleckchem, E1022) and TPA (12-O-tetradecanoylphorbol-13-acetate) (Sigma-Aldrich, P1585). The dorsal skin was initially treated with 100 µg DMBA applied topically twice at one-week intervals, followed by twice-weekly topical applications of 12.5 µg TPA. Tumor onset and progression were monitored daily by visual inspection and palpation. Mice were euthanized 33 weeks after treatment initiation. After euthanasia, normal and tumor-bearing dorsal skin samples were collected, fixed in 4% paraformaldehyde (PFA), and embedded in paraffin. All procedures were conducted in accordance with the European Community Council Directive 86/609/EEC and were approved by the CEINGE Ethical Committee for Animal Experimentation and the Italian Ministry of Health (D5A89.94, auth. 1102/2024-PR).

Intradermal ear injection was performed essentially as described in (98). SCC cells grown in DMEM supplemented with 10% FBS, were freshly infected with shRNA, and selected in puromycin. One week after infection, cells were labelled with PKH26 Red Lipid Dye (Sigma-Aldrich, MINI26-1KT, Sigma) prior to injection. A total of 200,000 cells were injected into each ear of four-week-old male NOD/SCID mice, with control cells injected into one ear and shRNA-expressing cells into the contralateral ear of the same animal. Five mice were used for each shRNA experiment. Equal number of injected cells was assessed the following day by immunofluorescence. The experiment was terminated after 45 days. All animal procedures were performed in accordance with Swiss regulations on laboratory animal use and were approved by the Veterinary Office of the Canton of Vaud.

### Tissue immunostaining

Formalin-fixed, paraffin-embedded human skin sections were stained with mouse monoclonal antibodies for p63 (Santa Cruz Biotechnology, 4A4) and rabbit polyclonal antibodies for p73 (Abcam, EPR5701) and counterstained using 4′,6-diamidino-2-phenylindole (DAPI; Thermo Fisher D1306). Human tissue microarray samples were previously described and consisted of normal skin (N=28), actinic keratosis (N=23), and SCC (N=24) (99). Tissues were imaged using a Microscope Leica DM5500 (Leica Microsystems) and corrected total cell fluorescence was calculated using ImageJ software as previously described (100).

Formalin-fixed, paraffin-embedded mouse skin tumors were sectioned at 5μm and processed for immunofluorescence using standard procedures. For antigen retrieval, sections were subjected to heat-induced epitope retrieval in 0.01 M citrate buffer (pH 6.0). Tissue was stained with anti-p73 antibodies, and counterstained using DAPI. Fluorescence images were acquired using an Axio Scan.Z1 microscope (Zeiss) at the Advanced Light Microscopy (ALM) Facility, CEINGE, and analyzed with ZEN 3.10 software (Zeiss).

### Real time RT-qPCR

Total RNA was extracted 48 hr after transfection from SCC cells and HK using TRIzol reagent (Thermo Fisher, 15596026). Gene expression levels were normalized to RPLP0. cDNA was synthesized with SuperScript™ VILO™ Master Mix (Thermo Fisher, 11755050) and target genes were quantified using ABI PRISM 7500 Real-Time PCR System (Applied Biosystems), using SYBR Select Master mix (Thermo Fisher, 4472918).

### RNA-seq

Total RNA was extracted with RNeasy Mini Kit (QIAGEN, 28104). Libraries were prepared using the NextSeq 500/550 output kit (Illumina), and sequence on a HiSeq 4000 System (Illumina).

Fastq files were aligned to the human genome assembly hg38 (NCBI Genome assembly GRCh38) using STAR (101) with default parameters. Quantification and differential gene expression analyses were performed using FeatureCounts and DESeq2, respectively. For visualization, bigWig files were generated using wigToBigWig (UCSC Genome Browser tools) and uploaded to the UCSC Genome Browser. Genes with a DESeq2 baseMean greater than 5 were included in downstream analyses. Differential expression (adjusted P value < 0.05) and principal component analysis were performed using DESeq2 based on read counts per gene (102). Biological processes were identified using Gene Ontology (GO) Biological Process terms via Metascape (103). Gene Set Enrichment Analysis (55) was performed using a dataset of primary SCC versus matched normal sun-exposed or perilesional skin (56) (Figure 3C) or using the Hallmark gene sets (104) (Figure 3E).

### ChIP-seq

Chromatin was cross-linked with 1% formaldehyde for 10 min at 37°C and quenched with 125 mM glycine for 5 min at room temperature. Cells were washed in cold PBS containing protease inhibitors and lysed (4×10⁶ cells/sample). Chromatin was sonicated (Bioruptor Plus; high power, 60 s ON/90 s OFF, 45 min, 4°C) to an average size of ∼300 bp, cleared by centrifugation, and diluted in ChIP dilution buffer. Immunoprecipitations were performed overnight at 4°C using 2.1 mg Dynabeads Protein A/G (1:1) and 4 μg anti-p63 (Abcam, EPR5701) or anti-p73 (Abcam, EP436Y) antibodies. Beads were washed sequentially with low-salt, high-salt, LiCl, and TE buffers, and chromatin was eluted for 20 min at room temperature. Cross-links were reversed overnight at 65°C in the presence of Proteinase K, and DNA was purified using QIAquick columns. Re-ChIP assays were performed by re-immunoprecipitating p63-bound chromatin with anti-p73 antibodies or IgG control. ChIP-qPCR was performed using SYBR Select Master Mix on an ABI 7500 system with locus-specific primers.

For ChIP-seq, libraries were prepared using the KAPA HiFi library amplification kit (Roche, KK2611) and sequenced on an Illumina HiSeq 2000. Reads were uniquely aligned to the human genome (hg38) using Bowtie2. Peak calling was performed with MACS2 using matched input as background. Approximately 48,000 p63 peaks (FE>5) and 27,900 p73 peaks were identified. A stringent p63 dataset (23,956 peaks; FDR<10⁻⁴⁰, FE>5) was selected for downstream analyses, of which 94% overlapped with p73 peaks (FDR<10⁻⁵, FE>5).

Peak intersections with cCREs and H3K27ac ChIP-seq datasets were performed using BEDTools. Peak–gene associations were assigned using GREAT (basal plus extension model; ±5 kb proximal, up to 100 kb distal). Motif enrichment analysis was conducted on the top 1,000 peaks using HOMER with the JASPAR 2022 database. Peak clustering and heatmap visualization were performed using ChAsE.

To identify cell type–specific p63/p73 binding sites, p63/p73 peaks in SCC cells were compared with published p63 ChIP-seq datasets from newborn and adult human keratinocytes and oral SCC (OSCC). A 250 bp window centered on the summits of high-confidence p63/p73 peaks in SCC cells (13,456 regions; p63 FDR<10⁻⁶⁰, FE>20; p73 FDR<10⁻^20^, FE>10) was intersected with p63 peaks from OSCC cells (21,461 regions; FE>10; GSE104145). Binding regions common to SCC and OSCC (5,950 regions, ≥250 bp) were removed from regions shared with newborn keratinocytes (30,106 regions; GSE56674) and adult keratinocytes (46,716 regions; GSE59824), resulting in 822 bona fide SCC-specific p63/p73 binding sites. To identify cutaneous-specific binding sites, p63/p73 peaks shared between SCC cells, newborn keratinocytes, and adult keratinocytes (21,236 regions; FDR<10⁻⁴⁰, FE>5; GSE59824) were intersected and regions overlapping p63 peaks in OSCC (39,582 regions; GSE104145) were excluded. This analysis identified 2,785 *bona fide* cutaneous-specific p63/p73 binding sites.

### Cell cycle analysis and apoptosis

To evaluate DNA synthesis, cells were labeled with BrdU Labeling Reagent (Thermo Fisher, 000103) for 2 hr, and subsequently fixed with 4% paraformaldehyde. After fixation, cells were permeabilized with NP-40 0.1%, and the DNA was denatured with 50 mM NaOH. BrdU was detected with mouse monoclonal antibodies (Developmental Studies Hybridoma Bank, The University of Iowa, G3G4) and goat-anti mouse Alexa Fluor 594 (Thermo Fisher, A-11005). DNA was counterstained with DAPI (Thermo Fisher, D1306). For cell cycle analysis, cells were trypsinized, washed three times with ice-cold PBS, and fully resuspended to avoid clumping. After a 5-min incubation on ice, PBS was removed and cells were resuspended in 300 μl ice-cold PBS. While vortexing gently, 700 μl of ice-cold 100% ethanol was added dropwise, followed by additional pipetting to ensure complete dispersion.

Fixed cells were passed through a 40-μm strainer and incubated at 4°C for 15 min in a fresh staining solution (PBS/EDTA supplemented with 0.1% Triton X-100, RNase A, and propidium iodide (Sigma-Aldrich, P4864)). Cell-cycle profiles were acquired by flow cytometry and quantified using ModFit LT software (Verity Software House).

To obtain more accurate cell-cycle phase identification, EdU incorporation was combined with DNA-content measurement using the Click-iT EdU Alexa Fluor 488 Flow Cytometry Assay Kit (Thermo Fisher 35002) and DRAQ5 (Thermo Fisher 62251). EdU was added to the culture medium at a final concentration of 10 μM for 2 h. Cells were then washed in PBS containing 1% BSA and fixed in the Click-iT fixative solution for 15 min at room temperature. Following an additional wash with PBS/1% BSA, cells were permeabilized in the kit’s saponin-based permeabilization buffer and labeled according to the manufacturer’s instructions. After DNA staining with DRAQ5, samples were acquired on a FACSCanto II flow cytometer (BD Biosciences) and analyzed using BD FACSDiva software. FACS analysis with Annexin V-FITC was performed in SCC cells 48hr upon transfection, or in SCC cells upon 2hr from UV treatment (1500 mJ/cm^2^) as positive control using a Stratalinker 1800 (Stratagene, La Jolla, CA, USA).

### Proximity ligation assay (PLA)

SCC cells were fixed 48 hr after siRNA transfection for 30’ at room temperature with 2% Paraformaldehyde. Proximity ligation assays (PLA) were performed using Duolink PLA (Sigma-Aldrich; DUO92101) according to the protocol provided by the manufacturer. Primary antibody used for PLA were diluted as follow: p63 1:20 (Abcam, AB735); p73 (Abcam, AB40658). Imaging was performed using confocal microscopy (ZEISS LSM980) and PLA spots were quantified using ZEN 3.10 software.

For the transfection SCC were treated the day after seeding with specific Stealth siRNA targeting p63 or p73 using Polyplus-transfection reagent INTERFERin® (Sartorius, 101000028) diluted 5:1000 v/v in serum-free K-SFM). Six hr later medium was changed with supplemented K-SFM and the experiment ended 48 hr later.

### Immunoprecipitation, immunoblotting and ELISA assay

SCC cells were lysed with an intermediate stringency buffer (Tris HCl 50 mM pH 8, NaCl 120 mM, NP-40 0.5%), supplemented with protease and phosphatase inhibitors (Roche-Merck 11697498001, Sigma Aldrich 4906837001) and incubated for 10 min on ice. After centrifugation at maximum speed, cleared protein extracts were incubated with primary antibodies pre-coupled to Dynabeads® Protein A and G (Thermo Fisher, 10001D, 10003D). Following 2hr incubation with gentle rotation, immune complexes were washed four times and eluted by boiling in Laemmli sample buffer. For immunoblotting, cells were lysed directly in Laemmli sample buffer, and proteins were resolved by SDS-PAGE, transferred to Immobilon-P PVDF membranes (Millipore-Merck, IPVH00010), probed with the indicated primary antibodies, and detected by chemiluminescence (Bio-Rad, 1705061). ELISA measurements were performed on culture supernatants collected 48, 72, and 96 hr after p63 depletion or negative control transfection, using the Human Amphiregulin ELISA Kit (Sigma Aldrich, RAB0019) following the manufacturer’s protocol.

## Data availability

Sequencing data have been deposited at GEO and are accessible through GEO Series accession number GSE235467 (RNA-Seq GSE235459; ChIP-Seq GSE235461).

## Acknowledgements

We gratefully acknowledge the financial support of the Italian Association for Cancer Research (AIRC IG 17079 and IG25116) to C.M., COINOR, University of Naples Federico II to D.A. (17-CSP-UNINA-060), the National Institutes of Health to M.K. (1R01CA154715), the Fondazione Gabriella Fabbrocini award to CM. We would like to thank Teresa Mattiello for performing preliminary experiments and Dania Al-Labban for help with xenografting. In addition, we thank the CIRO Project: “Campania Imaging Infrastructure for Research in Oncology” and the Advanced Light Microscopy Facility, the FACS facility, and the Animal Facility at CEINGE, and the Computational Biology Core at the Department of Biology of University of Naples Federico II.

## Author contributions

D.A. and C.M. conceived the study, interpreted most of the experiments, prepared the figures, and wrote the manuscript. D.A. and M. Ferniani performed the ChIP-seq and RNA-seq experiments, immunostaining of human tissues, and associated data analyses. M. Ferniani and C.R. carried out the depletion experiments and qPCR analyses. M.Ferniani performed clonal growth assays and tumorsphere assays. J.Q. prepared the ChIP-seq library and performed sequencing, and H.Z. supervised and performed the first analyses of the ChIP-seq datasets. M. Franciosi conducted the proximity ligation assays, cell-cycle analyses, and co-immunoprecipitation experiments. L.D. and M. Franciosi generated and analyzed the mouse carcinogenesis model. S.P. validated selected key experiments. A.S. provided SCC cell lines. M.K. provided normal skin, AK, and SCC tissue arrays. G.P.D. provided SCC cell lines and contributed expert input and interpretation of the tumor-injection experiments performed in his laboratory. All authors discussed the results and commented on the manuscript.

## References

1. Winge, M.C.G., Kellman, L.N., Guo, K., Tang, J.Y., Swetter, S.M., Aasi, S.Z., Sarin, K.Y., Chang, A.L.S. and Khavari, P.A. (2023) Advances in cutaneous squamous cell carcinoma. Nature reviews. Cancer, 23, 430–449.

2. Martincorena, I., Roshan, A., Gerstung, M., Ellis, P., Van Loo, P., McLaren, S., Wedge, D.C., Fullam, A., Alexandrov, L.B., Tubio, J.M., et al. (2015) Tumor evolution. High burden and pervasive positive selection of somatic mutations in normal human skin. Science, 348, 880–886.

3. Chitsazzadeh, V., Coarfa, C., Drummond, J.A., Nguyen, T., Joseph, A., Chilukuri, S., Charpiot, E., Adelmann, C.H., Ching, G., Nguyen, T.N. et al. (2016) Cross-species identification of genomic drivers of squamous cell carcinoma development across preneoplastic intermediates. Nat Commun, 7, 12601.

4. Chang, D. and Shain, A.H. (2021) The landscape of driver mutations in cutaneous squamous cell carcinoma. NPJ Genom Med, 6, 61.

5. Paull, E.O., Aytes, A., Jones, S.J., Subramaniam, P.S., Giorgi, F.M., Douglass, E.F., Tagore, S., Chu, B., Vasciaveo, A., Zheng, S. et al. (2021) A modular master regulator landscape controls cancer transcriptional identity. Cell, 184, 334–351 e320.

6. Osterburg, C. and Dotsch, V. (2022) Structural diversity of p63 and p73 isoforms. Cell death and differentiation, 29, 921–937.

7. Saladi, S.V., Ross, K., Karaayvaz, M., Tata, P.R., Mou, H., Rajagopal, J., Ramaswamy, S. and Ellisen, L.W. (2017) ACTL6A Is Co-Amplified with p63 in Squamous Cell Carcinoma to Drive YAP Activation, Regenerative Proliferation, and Poor Prognosis. Cancer cell, 31, 35–49.

8. Fisher, M.L., Balinth, S. and Mills, A.A. (2023) DeltaNp63alpha in cancer: importance and therapeutic opportunities. Trends Cell Biol, 33, 280–292.

9. Prieto-Garcia, C., Hartmann, O., Reissland, M., Braun, F., Fischer, T., Walz, S., Schulein-Volk, C., Eilers, U., Ade, C.P., Calzado, M.A. et al. (2020) Maintaining protein stability of Np63 via USP28 is required by squamous cancer cells. EMBO molecular medicine, 12, e11101.

10. Campbell, J.D., Yau, C., Bowlby, R., Liu, Y., Brennan, K., Fan, H., Taylor, A.M., Wang, C., Walter, V., Akbani, R. et al. (2018) Genomic, Pathway Network, and Immunologic Features Distinguishing Squamous Carcinomas. Cell reports, 23, 194–212 e196.

11. Satpathy, S., Krug, K., Jean Beltran, P.M., Savage, S.R., Petralia, F., Kumar-Sinha, C., Dou, Y., Reva, B., Kane, M.H., Avanessian, S.C. et al. (2021) A proteogenomic portrait of lung squamous cell carcinoma. Cell, 184, 4348–4371 e4340.

12. Zhang, Y., Karagiannis, D., Liu, H., Lin, M., Fang, Y., Jiang, M., Chen, X., Suresh, S., Huang, H., She, J. et al. (2024) Epigenetic regulation of p63 blocks squamous-to-neuroendocrine transdifferentiation in esophageal development and malignancy. Sci Adv, 10, eadq0479.

13. Fisher, M.L., Balinth, S., Hwangbo, Y., Wu, C., Ballon, C., Wilkinson, J.E., Goldberg, G.L. and Mills, A.A. (2021) BRD4 Regulates Transcription Factor DeltaNp63alpha to Drive a Cancer Stem Cell Phenotype in Squamous Cell Carcinomas. Cancer research, 81, 6246–6258.

14. Abraham, C.G., Ludwig, M.P., Andrysik, Z., Pandey, A., Joshi, M., Galbraith, M.D., Sullivan, K.D. and Espinosa, J.M. (2018) DeltaNp63alpha Suppresses TGFB2 Expression and RHOA Activity to Drive Cell Proliferation in Squamous Cell Carcinomas. Cell reports, 24, 3224–3236.

15. Kajiwara, C., Fumoto, K., Kimura, H., Nojima, S., Asano, K., Odagiri, K., Yamasaki, M., Hikita, H., Takehara, T., Doki, Y. et al. (2018) p63-Dependent Dickkopf3 Expression Promotes Esophageal Cancer Cell Proliferation via CKAP4. Cancer research, 78, 6107–6120.

16. Ramsey, M.R., He, L., Forster, N., Ory, B. and Ellisen, L.W. (2011) Physical association of HDAC1 and HDAC2 with p63 mediates transcriptional repression and tumor maintenance in squamous cell carcinoma. Cancer research, 71, 4373–4379.

17. Rocco, J.W., Leong, C.O., Kuperwasser, N., DeYoung, M.P. and Ellisen, L.W. (2006) p63 mediates survival in squamous cell carcinoma by suppression of p73-dependent apoptosis. Cancer cell, 9, 45–56.

18. Hsieh, M.H., Choe, J.H., Gadhvi, J., Kim, Y.J., Arguez, M.A., Palmer, M., Gerold, H., Nowak, C., Do, H., Mazambani, S. et al. (2019) p63 and SOX2 Dictate Glucose Reliance and Metabolic Vulnerabilities in Squamous Cell Carcinomas. Cell reports, 28, 1860–1878 e1869.

19. Dong, J., Li, J., Li, Y., Ma, Z., Yu, Y. and Wang, C.Y. (2021) Transcriptional super-enhancers control cancer stemness and metastasis genes in squamous cell carcinoma. Nat Commun, 12, 3974.

20. Watanabe, H., Ma, Q., Peng, S., Adelmant, G., Swain, D., Song, W., Fox, C., Francis, J.M., Pedamallu, C.S., DeLuca, D.S. et al. (2014) SOX2 and p63 colocalize at genetic loci in squamous cell carcinomas. The Journal of clinical investigation, 124, 1636–1645.

21. Li, L.Y., Yang, Q., Jiang, Y.Y., Yang, W., Jiang, Y., Li, X., Hazawa, M., Zhou, B., Huang, G.W., Xu, X.E. et al. (2021) Interplay and cooperation between SREBF1 and master transcription factors regulate lipid metabolism and tumor-promoting pathways in squamous cancer. Nat Commun, 12, 4362.

22. Keyes, W.M., Pecoraro, M., Aranda, V., Vernersson-Lindahl, E., Li, W., Vogel, H., Guo, X., Garcia, E.L., Michurina, T.V., Enikolopov, G. et al. (2011) DeltaNp63alpha is an oncogene that targets chromatin remodeler Lsh to drive skin stem cell proliferation and tumorigenesis. Cell stem cell, 8, 164–176.

23. Westfall, M.D., Mays, D.J., Sniezek, J.C. and Pietenpol, J.A. (2003) The Delta Np63 alpha phosphoprotein binds the p21 and 14-3-3 sigma promoters in vivo and has transcriptional repressor activity that is reduced by Hay-Wells syndrome-derived mutations. Molecular and cellular biology, 23, 2264–2276.

24. DeYoung, M.P., Johannessen, C.M., Leong, C.O., Faquin, W., Rocco, J.W. and Ellisen, L.W. (2006) Tumor-specific p73 up-regulation mediates p63 dependence in squamous cell carcinoma. Cancer research, 66, 9362–9368.

25. Truong, A.B., Kretz, M., Ridky, T.W., Kimmel, R. and Khavari, P.A. (2006) p63 regulates proliferation and differentiation of developmentally mature keratinocytes. Genes & development, 20, 3185–3197.

26. Antonini, D., Russo, M.T., De Rosa, L., Gorrese, M., Del Vecchio, L. and Missero, C. (2010) Transcriptional repression of miR-34 family contributes to p63-mediated cell cycle progression in epidermal cells. The Journal of investigative dermatology, 130, 1249–1257.

27. Gallant-Behm, C.L. and Espinosa, J.M. (2013) DeltaNp63alpha utilizes multiple mechanisms to repress transcription in squamous cell carcinoma cells. Cell cycle, 12, 409–416.

28. Riege, K., Kretzmer, H., Sahm, A., McDade, S.S., Hoffmann, S. and Fischer, M. (2020) Dissecting the DNA binding landscape and gene regulatory network of p63 and p53. Elife, 9.

29. Bao, X., Rubin, A.J., Qu, K., Zhang, J., Giresi, P.G., Chang, H.Y. and Khavari, P.A. (2015) A novel ATAC-seq approach reveals lineage-specific reinforcement of the open chromatin landscape via cooperation between BAF and p63. Genome Biol, 16, 284.

30. Ramsey, M.R., Wilson, C., Ory, B., Rothenberg, S.M., Faquin, W., Mills, A.A. and Ellisen, L.W. (2013) FGFR2 signaling underlies p63 oncogenic function in squamous cell carcinoma. The Journal of clinical investigation, 123, 3525–3538.

31. Devos, M., Gilbert, B., Denecker, G., Leurs, K., Mc Guire, C., Lemeire, K., Hochepied, T., Vuylsteke, M., Lambert, J., Van Den Broecke, C., et al. (2017) Elevated DeltaNp63alpha Levels Facilitate Epidermal and Biliary Oncogenic Transformation. The Journal of investigative dermatology, 137, 494–505.

32. Sastre-Perona, A., Hoang-Phou, S., Leitner, M.C., Okuniewska, M., Meehan, S. and Schober, M. (2019) De Novo PITX1 Expression Controls Bi-Stable Transcriptional Circuits to Govern Self-Renewal and Differentiation in Squamous Cell Carcinoma. Cell stem cell, 24, 390–404 e398.

33. Song, Q., Yang, Y., Jiang, D., Qin, Z., Xu, C., Wang, H., Huang, J., Chen, L., Luo, R., Zhang, X. et al. (2022) Proteomic analysis reveals key differences between squamous cell carcinomas and adenocarcinomas across multiple tissues. Nat Commun, 13, 4167.

34. Johnson, J., Lagowski, J., Sundberg, A., Lawson, S., Liu, Y. and Kulesz-Martin, M. (2007) p73 loss triggers conversion to squamous cell carcinoma reversible upon reconstitution with TAp73alpha. Cancer research, 67, 7723–7730.

35. Lu, H., Yang, X., Duggal, P., Allen, C.T., Yan, B., Cohen, J., Nottingham, L., Romano, R.A., Sinha, S., King, K.E. et al. (2011) TNF-alpha promotes c-REL/DeltaNp63alpha interaction and TAp73 dissociation from key genes that mediate growth arrest and apoptosis in head and neck cancer. Cancer research, 71, 6867–6877.

36. Coutandin, D., Lohr, F., Niesen, F.H., Ikeya, T., Weber, T.A., Schafer, B., Zielonka, E.M., Bullock, A.N., Yang, A., Guntert, P. et al. (2009) Conformational stability and activity of p73 require a second helix in the tetramerization domain. Cell death and differentiation, 16, 1582–1589.

37. Davison, T.S., Vagner, C., Kaghad, M., Ayed, A., Caput, D. and Arrowsmith, C.H. (1999) p73 and p63 are homotetramers capable of weak heterotypic interactions with each other but not with p53. The Journal of biological chemistry, 274, 18709–18714.

38. Gebel, J., Luh, L.M., Coutandin, D., Osterburg, C., Lohr, F., Schafer, B., Frombach, A.S., Sumyk, M., Buchner, L., Krojer, T. et al. (2016) Mechanism of TAp73 inhibition by DeltaNp63 and structural basis of p63/p73 hetero-tetramerization. Cell death and differentiation, 23, 1930–1940.

39. Strubel, A., Munick, P., Hartmann, O., Chaikuad, A., Dreier, B., Schaefer, J.V., Gebel, J., Osterburg, C., Tuppi, M., Schafer, B. et al. (2023) DARPins detect the formation of hetero-tetramers of p63 and p73 in epithelial tissues and in squamous cell carcinoma. Cell death & disease, 14, 674.

40. Cancer Genome Atlas Research, N., Analysis Working Group: Asan, U., Agency, B.C.C., Brigham, Women’s, H., Broad, I., Brown, U., Case Western Reserve, U., Dana-Farber Cancer, I., Duke, U., et al. (2017) Integrated genomic characterization of oesophageal carcinoma. Nature, 541, 169–175.

41. Cancer Genome Atlas Research, N., Albert Einstein College of, M., Analytical Biological, S., Barretos Cancer, H., Baylor College of, M., Beckman Research Institute of City of, H., Buck Institute for Research on, A., Canada’s Michael Smith Genome Sciences, C., Harvard Medical, S., Helen, F.G.C.C. et al. (2017) Integrated genomic and molecular characterization of cervical cancer. Nature, 543, 378–384.

42. Cancer Genome Atlas, N. (2015) Comprehensive genomic characterization of head and neck squamous cell carcinomas. Nature, 517, 576–582.

43. Joseph, S.R., Gaffney, D., Barry, R., Hu, L., Banushi, B., Wells, J.W., Lambie, D., Strutton, G., Porceddu, S.V., Burmeister, B. et al. (2019) An Ex Vivo Human Tumor Assay Shows Distinct Patterns of EGFR Trafficking in Squamous Cell Carcinoma Correlating to Therapeutic Outcomes. The Journal of investigative dermatology, 139, 213–223.

44. Canueto, J., Cardenoso, E., Garcia, J.L., Santos-Briz, A., Castellanos-Martin, A., Fernandez-Lopez, E., Blanco Gomez, A., Perez-Losada, J. and Roman-Curto, C. (2017) Epidermal growth factor receptor expression is associated with poor outcome in cutaneous squamous cell carcinoma. The British journal of dermatology, 176, 1279–1287.

45. Rittie, L., Kansra, S., Stoll, S.W., Li, Y., Gudjonsson, J.E., Shao, Y., Michael, L.E., Fisher, G.J., Johnson, T.M. and Elder, J.T. (2007) Differential ErbB1 signaling in squamous cell versus basal cell carcinoma of the skin. The American journal of pathology, 170, 2089–2099.

46. Stoll, S.W., Johnson, J.L., Bhasin, A., Johnston, A., Gudjonsson, J.E., Rittie, L. and Elder, J.T. (2010) Metalloproteinase-mediated, context-dependent function of amphiregulin and HB-EGF in human keratinocytes and skin. The Journal of investigative dermatology, 130, 295–304.

47. Stoll, S.W., Stuart, P.E., Swindell, W.R., Tsoi, L.C., Li, B., Gandarillas, A., Lambert, S., Johnston, A., Nair, R.P. and Elder, J.T. (2016) The EGF receptor ligand amphiregulin controls cell division via FoxM1. Oncogene, 35, 2075–2086.

48. Marshall, C.B., Beeler, J.S., Lehmann, B.D., Gonzalez-Ericsson, P., Sanchez, V., Sanders, M.E., Boyd, K.L. and Pietenpol, J.A. (2021) Tissue-specific expression of p73 and p63 isoforms in human tissues. Cell death & disease, 12, 745.

49. Uhlen, M., Zhang, C., Lee, S., Sjostedt, E., Fagerberg, L., Bidkhori, G., Benfeitas, R., Arif, M., Liu, Z., Edfors, F. et al. (2017) A pathology atlas of the human cancer transcriptome. Science, 357.

50. Marshall, C.B., Mays, D.J., Beeler, J.S., Rosenbluth, J.M., Boyd, K.L., Santos Guasch, G.L., Shaver, T.M., Tang, L.J., Liu, Q., Shyr, Y. et al. (2016) p73 Is Required for Multiciliogenesis and Regulates the Foxj1-Associated Gene Network. Cell reports, 14, 2289–2300.

51. Nemajerova, A., Kramer, D., Siller, S.S., Herr, C., Shomroni, O., Pena, T., Gallinas Suazo, C., Glaser, K., Wildung, M., Steffen, H. et al. (2016) TAp73 is a central transcriptional regulator of airway multiciliogenesis. Genes & development, 30, 1300–1312.

52. Consortium, E.P., Moore, J.E., Purcaro, M.J., Pratt, H.E., Epstein, C.B., Shoresh, N., Adrian, J., Kawli, T., Davis, C.A., Dobin, A. et al. (2022) Author Correction: Expanded encyclopaedias of DNA elements in the human and mouse genomes. Nature, 605, E3.

53. Kouwenhoven, E.N., Oti, M., Niehues, H., van Heeringen, S.J., Schalkwijk, J., Stunnenberg, H.G., van Bokhoven, H. and Zhou, H. (2015) Transcription factor p63 bookmarks and regulates dynamic enhancers during epidermal differentiation. EMBO Rep, 16, 863–878.

54. McDade, S.S., Henry, A.E., Pivato, G.P., Kozarewa, I., Mitsopoulos, C., Fenwick, K., Assiotis, I., Hakas, J., Zvelebil, M., Orr, N. et al. (2012) Genome-wide analysis of p63 binding sites identifies AP-2 factors as co-regulators of epidermal differentiation. Nucleic acids research, 40, 7190–7206.

55. Subramanian, A., Tamayo, P., Mootha, V.K., Mukherjee, S., Ebert, B.L., Gillette, M.A., Paulovich, A., Pomeroy, S.L., Golub, T.R., Lander, E.S. et al. (2005) Gene set enrichment analysis: a knowledge-based approach for interpreting genome-wide expression profiles. Proceedings of the National Academy of Sciences of the United States of America, 102, 15545–15550.

56. Bailey, P., Ridgway, R.A., Cammareri, P., Treanor-Taylor, M., Bailey, U.M., Schoenherr, C., Bone, M., Schreyer, D., Purdie, K., Thomson, J. et al. (2023) Driver gene combinations dictate cutaneous squamous cell carcinoma disease continuum progression. Nat Commun, 14, 5211.

57. Joerger, A.C., Rajagopalan, S., Natan, E., Veprintsev, D.B., Robinson, C.V. and Fersht, A.R. (2009) Structural evolution of p53, p63, and p73: implication for heterotetramer formation. Proceedings of the National Academy of Sciences of the United States of America, 106, 17705–17710.

58. Della Gatta, G., Bansal, M., Ambesi-Impiombato, A., Antonini, D., Missero, C. and di Bernardo, D. (2008) Direct targets of the TRP63 transcription factor revealed by a combination of gene expression profiling and reverse engineering. Genome research, 18, 939–948.

59. Sethi, I., Gluck, C., Zhou, H., Buck, M.J. and Sinha, S. (2017) Evolutionary re-wiring of p63 and the epigenomic regulatory landscape in keratinocytes and its potential implications on species-specific gene expression and phenotypes. Nucleic acids research, 45, 8208–8224.

60. Li, L., Wang, Y., Torkelson, J.L., Shankar, G., Pattison, J.M., Zhen, H.H., Fang, F., Duren, Z., Xin, J., Gaddam, S. et al. (2019) TFAP2C– and p63-Dependent Networks Sequentially Rearrange Chromatin Landscapes to Drive Human Epidermal Lineage Commitment. Cell stem cell, 24, 271–284 e278.

61. Santos-Pereira, J.M., Gallardo-Fuentes, L., Neto, A., Acemel, R.D. and Tena, J.J. (2019) Pioneer and repressive functions of p63 during zebrafish embryonic ectoderm specification. Nat Commun, 10, 3049.

62. Glathar, A.R., Oyelakin, A., Nayak, K.B., Sosa, J., Romano, R.A. and Sinha, S. (2023) A Systemic and Integrated Analysis of p63-Driven Regulatory Networks in Mouse Oral Squamous Cell Carcinoma. Cancers (Basel), 15.

63. Jiang, Y.Y., Jiang, Y., Li, C.Q., Zhang, Y., Dakle, P., Kaur, H., Deng, J.W., Lin, R.Y., Han, L., Xie, J.J. et al. (2020) TP63, SOX2, and KLF5 Establish a Core Regulatory Circuitry That Controls Epigenetic and Transcription Patterns in Esophageal Squamous Cell Carcinoma Cell Lines. Gastroenterology, 159, 1311–1327 e1319.

64. Gunschmann, C., Stachelscheid, H., Akyuz, M.D., Schmitz, A., Missero, C., Bruning, J.C. and Niessen, C.M. (2013) Insulin/IGF-1 controls epidermal morphogenesis via regulation of FoxO-mediated p63 inhibition. Dev Cell, 26, 176–187.

65. Kurinna, S., Seltmann, K., Bachmann, A.L., Schwendimann, A., Thiagarajan, L., Hennig, P., Beer, H.D., Mollo, M.R., Missero, C. and Werner, S. (2021) Interaction of the NRF2 and p63 transcription factors promotes keratinocyte proliferation in the epidermis. Nucleic acids research, 49, 3748–3763.

66. Consortium, E.P., Moore, J.E., Purcaro, M.J., Pratt, H.E., Epstein, C.B., Shoresh, N., Adrian, J., Kawli, T., Davis, C.A., Dobin, A. et al. (2020) Expanded encyclopaedias of DNA elements in the human and mouse genomes. Nature, 583, 699–710.

67. Goruppi, S., Clocchiatti, A., Bottoni, G., Di Cicco, E., Ma, M., Tassone, B., Neel, V., Demehri, S., Simon, C. and Paolo Dotto, G. (2023) The ULK3 kinase is a determinant of keratinocyte self-renewal and tumorigenesis targeting the arginine methylome. Nat Commun, 14, 887.

68. Bakkers, J., Hild, M., Kramer, C., Furutani-Seiki, M. and Hammerschmidt, M. (2002) Zebrafish DeltaNp63 is a direct target of Bmp signaling and encodes a transcriptional repressor blocking neural specification in the ventral ectoderm. Dev Cell, 2, 617–627.

69. De Rosa, L., Antonini, D., Ferone, G., Russo, M.T., Yu, P.B., Han, R. and Missero, C. (2009) p63 Suppresses non-epidermal lineage markers in a bone morphogenetic protein-dependent manner via repression of Smad7. The Journal of biological chemistry, 284, 30574–30582.

70. Kouwenhoven, E.N., van Heeringen, S.J., Tena, J.J., Oti, M., Dutilh, B.E., Alonso, M.E., de la Calle-Mustienes, E., Smeenk, L., Rinne, T., Parsaulian, L., et al. (2010) Genome-wide profiling of p63 DNA-binding sites identifies an element that regulates gene expression during limb development in the 7q21 SHFM1 locus. PLoS Genet, 6, e1001065.

71. Antonini, D., Rossi, B., Han, R., Minichiello, A., Di Palma, T., Corrado, M., Banfi, S., Zannini, M., Brissette, J.L. and Missero, C. (2006) An autoregulatory loop directs the tissue-specific expression of p63 through a long-range evolutionarily conserved enhancer. Molecular and cellular biology, 26, 3308–3318.

72. Antonini, D., Sirico, A., Aberdam, E., Ambrosio, R., Campanile, C., Fagoonee, S., Altruda, F., Aberdam, D., Brissette, J.L. and Missero, C. (2015) A composite enhancer regulates p63 gene expression in epidermal morphogenesis and in keratinocyte differentiation by multiple mechanisms. Nucleic acids research, 43, 862–874.

73. Rodriguez-Colman, M.J., Dansen, T.B. and Burgering, B.M.T. (2024) FOXO transcription factors as mediators of stress adaptation. Nat Rev Mol Cell Biol, 25, 46–64.

74. Gonzalez-Cano, L., Fuertes-Alvarez, S., Robledinos-Anton, N., Bizy, A., Villena-Cortes, A., Farinas, I., Marques, M.M. and Marin, M.C. (2016) p73 is required for ependymal cell maturation and neurogenic SVZ cytoarchitecture. Dev Neurobiol, 76, 730–747.

75. Sigismund, S., Avanzato, D. and Lanzetti, L. (2018) Emerging functions of the EGFR in cancer. Mol Oncol, 12, 3–20.

76. Stoll, S.W., Johnson, J.L., Li, Y., Rittie, L. and Elder, J.T. (2010) Amphiregulin carboxy-terminal domain is required for autocrine keratinocyte growth. The Journal of investigative dermatology, 130, 2031–2040.

77. Lambert, A.W., Fiore, C., Chutake, Y., Verhaar, E.R., Strasser, P.C., Chen, M.W., Farouq, D., Das, S., Li, X., Eaton, E.N. et al. (2022) DeltaNp63/p73 drive metastatic colonization by controlling a regenerative epithelial stem cell program in quasi-mesenchymal cancer stem cells. Dev Cell, 57, 2714–2730 e2718.

78. Mani, S.A., Guo, W., Liao, M.J., Eaton, E.N., Ayyanan, A., Zhou, A.Y., Brooks, M., Reinhard, F., Zhang, C.C., Shipitsin, M. et al. (2008) The epithelial-mesenchymal transition generates cells with properties of stem cells. Cell, 133, 704–715.

79. Logotheti, S., Pavlopoulou, A., Marquardt, S., Takan, I., Georgakilas, A.G. and Stiewe, T. (2022) p73 isoforms meet evolution of metastasis. Cancer Metastasis Rev, 41, 853–869.

80. McDonald, E.R., 3rd, de Weck, A., Schlabach, M.R., Billy, E., Mavrakis, K.J., Hoffman, G.R., Belur, D., Castelletti, D., Frias, E., Gampa, K. et al. (2017) Project DRIVE: A Compendium of Cancer Dependencies and Synthetic Lethal Relationships Uncovered by Large-Scale, Deep RNAi Screening. Cell, 170, 577–592 e510.

81. Kogashiwa, Y., Inoue, H., Kuba, K., Araki, R., Yasuda, M., Nakahira, M. and Sugasawa, M. (2018) Prognostic role of epiregulin/amphiregulin expression in recurrent/metastatic head and neck cancer treated with cetuximab. Head Neck, 40, 2424–2431.

82. Hsieh, M.J., Chen, Y.H., Lee, I.N., Huang, C., Ku, Y.J. and Chen, J.C. (2019) Secreted amphiregulin promotes vincristine resistance in oral squamous cell carcinoma. Int J Oncol, 55, 949–959.

83. Seligmann, J.F., Elliott, F., Richman, S.D., Jacobs, B., Hemmings, G., Brown, S., Barrett, J.H., Tejpar, S., Quirke, P. and Seymour, M.T. (2016) Combined Epiregulin and Amphiregulin Expression Levels as a Predictive Biomarker for Panitumumab Therapy Benefit or Lack of Benefit in Patients With RAS Wild-Type Advanced Colorectal Cancer. JAMA Oncol, 2, 633–642.

84. Conticelli, D., Volante, M., Pietrantonio, F., Orru, C., Olivero, M., Nottegar, A., Borghi, F., Baiocchi, G.L., Crotti, G., Fumagalli Romario, U. et al. (2025) AREG and EREG Are Predictive Biomarkers of Response to EGFR Inhibition in Gastroesophageal Cancer. Cancer research, 85, 3111–3122.

85. Qu, X., Hamidi, H., Johnson, R.M., Sokol, E.S., Lin, E., Eng, C., Kim, T.W., Bendell, J., Sivakumar, S., Kaplan, B. et al. (2025) Ligand-activated EGFR/MAPK signaling but not PI3K, are key resistance mechanisms to EGFR-therapy in colorectal cancer. Nat Commun, 16, 4332.

86. Montaudie, H., Viotti, J., Combemale, P., Dutriaux, C., Dupin, N., Robert, C., Mortier, L., Kaphan, R., Duval-Modeste, A.B., Dalle, S. et al. (2020) Cetuximab is efficient and safe in patients with advanced cutaneous squamous cell carcinoma: a retrospective, multicentre study. Oncotarget, 11, 378–385.

87. Migden, M.R., Rischin, D., Schmults, C.D., Guminski, A., Hauschild, A., Lewis, K.D., Chung, C.H., Hernandez-Aya, L., Lim, A.M., Chang, A.L.S. et al. (2018) PD-1 Blockade with Cemiplimab in Advanced Cutaneous Squamous-Cell Carcinoma. The New England journal of medicine, 379, 341–351.

88. Sidaway, P. (2018) Cemiplimab effective in cutaneous SCC. Nat Rev Clin Oncol, 15, 472.

89. de Lima, P.O., Joseph, S., Panizza, B. and Simpson, F. (2020) Epidermal Growth Factor Receptor’s Function in Cutaneous Squamous Cell Carcinoma and Its Role as a Therapeutic Target in the Age of Immunotherapies. Curr Treat Options Oncol, 21, 9.

90. Becker, J.C., Gesierich, A.H., Leiter, U., Zimmer, L., Hassel, J.C., von Wasielewski, I., Ziemer, M., Fluck, M., Meier, F., Spillner, A.N., et al. (2025) Avelumab plus cetuximab in patients with unresectable stage III or IV cutaneous squamous cell carcinoma – clinical activity and safety results from AliCe, a single-arm, multicentre phase 2 DeCOG trial. The British journal of dermatology.

91. Hamdan, F.H. and Johnsen, S.A. (2018) DeltaNp63-dependent super enhancers define molecular identity in pancreatic cancer by an interconnected transcription factor network. Proceedings of the National Academy of Sciences of the United States of America, 115, E12343–E12352.

92. Somerville, T.D.D., Xu, Y., Miyabayashi, K., Tiriac, H., Cleary, C.R., Maia-Silva, D., Milazzo, J.P., Tuveson, D.A. and Vakoc, C.R. (2018) TP63-Mediated Enhancer Reprogramming Drives the Squamous Subtype of Pancreatic Ductal Adenocarcinoma. Cell reports, 25, 1741–1755 e1747.

93. Hur, S.K., Somerville, T.D.D., Wu, X.S., Maia-Silva, D., Demerdash, O.E., Tuveson, D.A., Notta, F. and Vakoc, C.R. (2023) p73 activates transcriptional signatures of basal lineage identity and ciliogenesis in pancreatic ductal adenocarcinoma. bioRxiv.

94. Ebert, M., Yokoyama, M., Kobrin, M.S., Friess, H., Lopez, M.E., Buchler, M.W., Johnson, G.R. and Korc, M. (1994) Induction and expression of amphiregulin in human pancreatic cancer. Cancer research, 54, 3959–3962.

95. Mucciolo, G., Araos Henriquez, J., Jihad, M., Pinto Teles, S., Manansala, J.S., Li, W., Ashworth, S., Lloyd, E.G., Cheng, P.S.W., Luo, W. et al. (2024) EGFR-activated myofibroblasts promote metastasis of pancreatic cancer. Cancer cell, 42, 101–118 e111.

96. Su, X., Lai, T., Tao, Y., Zhang, Y., Zhao, C., Zhou, J., Chen, E., Zhu, M., Zhang, S., Wang, B. et al. (2023) miR-33a-3p regulates METTL3-mediated AREG stability and alters EMT to inhibit pancreatic cancer invasion and metastasis. Sci Rep, 13, 13587.

97. Wang, L., Wang, L., Zhang, H., Lu, J., Zhang, Z., Wu, H. and Liang, Z. (2020) AREG mediates the epithelialmesenchymal transition in pancreatic cancer cells via the EGFR/ERK/NFkappaB signalling pathway. Oncol Rep, 43, 1558–1568.

98. Procopio, M.G., Laszlo, C., Al Labban, D., Kim, D.E., Bordignon, P., Jo, S.H., Goruppi, S., Menietti, E., Ostano, P., Ala, U., et al. (2015) Combined CSL and p53 downregulation promotes cancer-associated fibroblast activation. Nature cell biology, 17, 1193–1204.

99. Hill, N.T., Zhang, J., Leonard, M.K., Lee, M., Shamma, H.N. and Kadakia, M. (2015) 1alpha, 25-Dihydroxyvitamin D(3) and the vitamin D receptor regulates DeltaNp63alpha levels and keratinocyte proliferation. Cell death & disease, 6, e1781.

100. Burgess, A., Vigneron, S., Brioudes, E., Labbe, J.C., Lorca, T. and Castro, A. (2010) Loss of human Greatwall results in G2 arrest and multiple mitotic defects due to deregulation of the cyclin B-Cdc2/PP2A balance. Proceedings of the National Academy of Sciences of the United States of America, 107, 12564–12569.

101. Dobin, A., Davis, C.A., Schlesinger, F., Drenkow, J., Zaleski, C., Jha, S., Batut, P., Chaisson, M. and Gingeras, T.R. (2013) STAR: ultrafast universal RNA-seq aligner. Bioinformatics, 29, 15–21.

102. Love, M.I., Huber, W. and Anders, S. (2014) Moderated estimation of fold change and dispersion for RNA-seq data with DESeq2. Genome Biol, 15, 550.

103. Zhou, Y., Zhou, B., Pache, L., Chang, M., Khodabakhshi, A.H., Tanaseichuk, O., Benner, C. and Chanda, S.K. (2019) Metascape provides a biologist-oriented resource for the analysis of systems-level datasets. Nat Commun, 10, 1523.

104. Liberzon, A., Birger, C., Thorvaldsdottir, H., Ghandi, M., Mesirov, J.P. and Tamayo, P. (2015) The Molecular Signatures Database (MSigDB) hallmark gene set collection. Cell Syst, 1, 417–425.

