## Supplementary Figures for "p63 and p73 regulate convergent and factor-specific transcriptional programs in cutaneous squamous cell carcinoma"

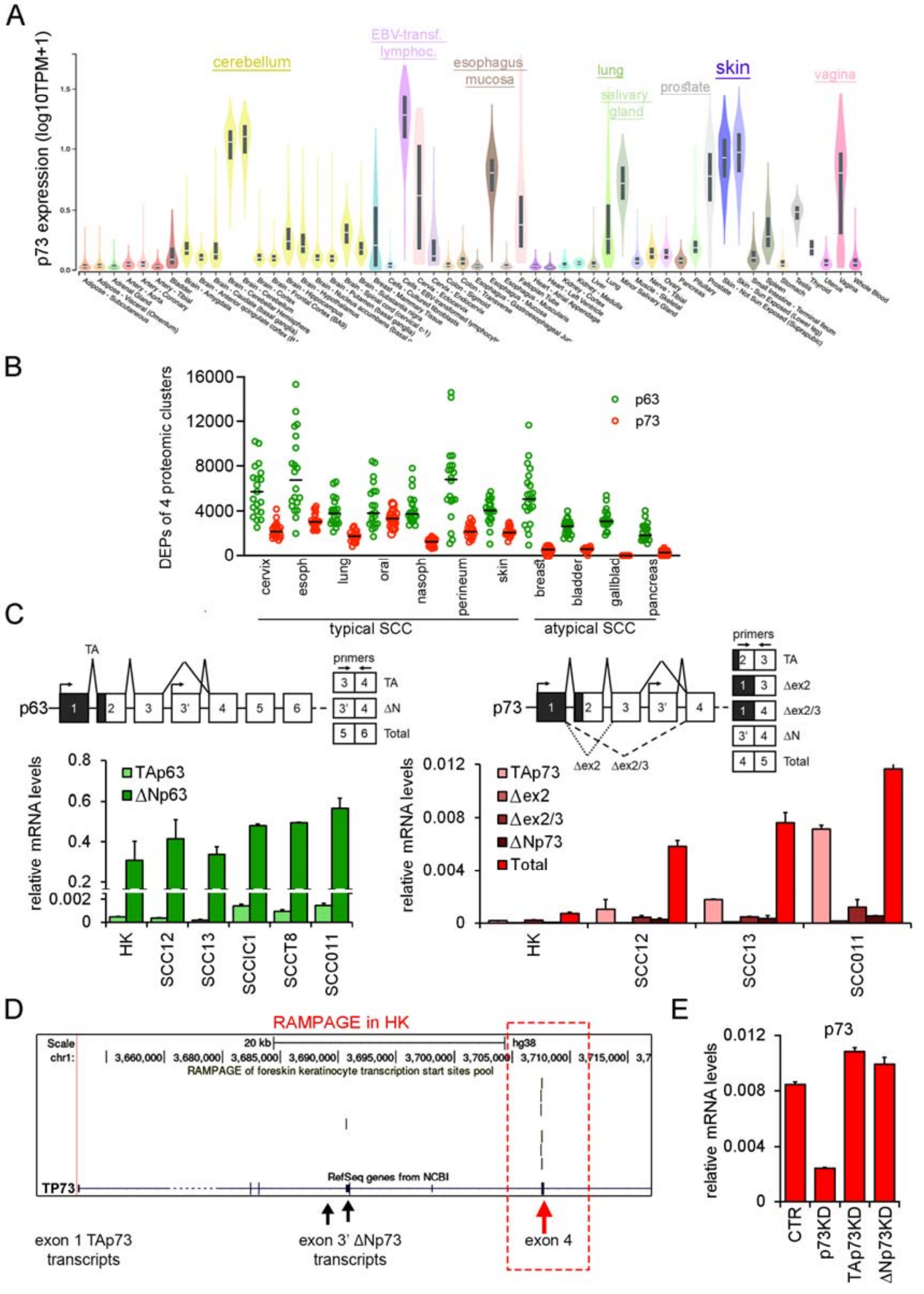


**Figure S1. p73 expression in skin and SCC, and isoform analysis of p63 and p73.**

(A) Bulk tissue gene expression of p73 (data from the GTEx project; https://gtexportal.org/home/). Gene expression values are shown as $\log_{10}$(TPM + 1). (B) p63 and p73 protein abundance in typical and atypical SCC, as indicated. Data were derived from a large-scale proteomic study (33). Each dot represents an individual SCC sample. Differentially expressed proteins (DEPs) for p63 and p73 were calculated using Kruskal-Wallis test with Benjamini–Hochberg correction (adjusted $p < 0.05$) across four clusters. across four proteomic clusters. In all tissues analyzed, p73 levels were significantly lower than p63. Moreover, p73 expression was significantly reduced in atypical compared with typical SCC. (C) Left: Schematic representation of the 5′ region of p63 showing exon structure, TSS (black arrows), TAp63 5'UTR (black), and primer locations (top). RT-qPCR analysis of TAp63 and ΔNp63 transcripts in the indicated cell types and (bottom). depletion Right: Schematic representation of the 5′ region of p73 as shown for p63 (top). RT-qPCR analysis of total p73 and transcript variants in the indicated cell types (bottom). (D) Genome browser view of the p73 5′ region showing the RAMPAGE signal in HK. (E) RT-qPCR analysis of total p73 mRNA levels in SCC13 cells transfected with the indicated siRNAs.

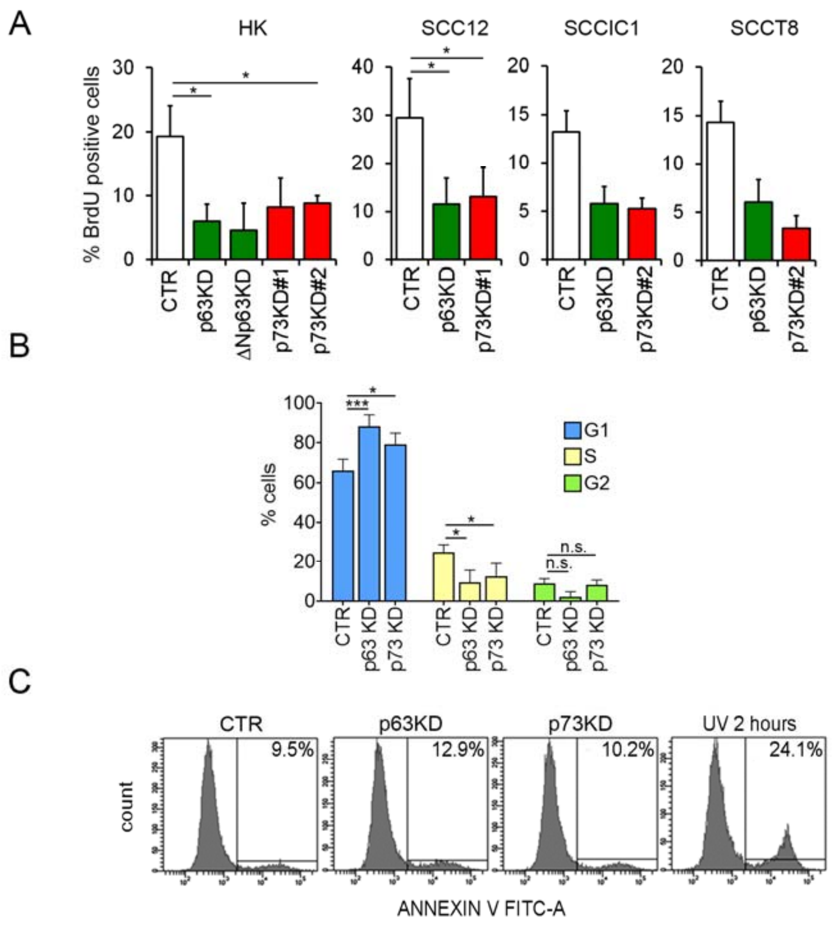


**Figure S2. Effects of p63 and p73 depletion on proliferation and apoptosis in SCC cells.** (A) Quantification of BrdU-positive cells relative to total nuclei in HK and in the indicated SCC cell lines. Data are shown as mean ± SD. Statistical analysis was performed using two-tailed Student's t-test. Replicate numbers: HK—CTR (n=5), p63KD (n=4), ΔNp63KD (n=2), p73KD#1 (n=2), p73KD#2 (n=2); SCC12—CTR (n=6), p63KD (n=6), p73KD#1 (n=3); SCCIC1 (n=1); SCCT8 (n=1). *P < 0.05. (B) Cell-cycle distribution of SCC cells after KD, assessed by propidium iodide staining and flow cytometry. Statistical analysis was performed using two-tailed Student's t-test. CTR (n=5), p63KD (n=3), p73KD (n=5). *P < 0.05; ***P < 0.0001. (C) Annexin V–FITC flow cytometric analysis of SCC13 cells after p63 or p73 KD, or UV treatment (2 h). Percentages of apoptotic cells are indicated.

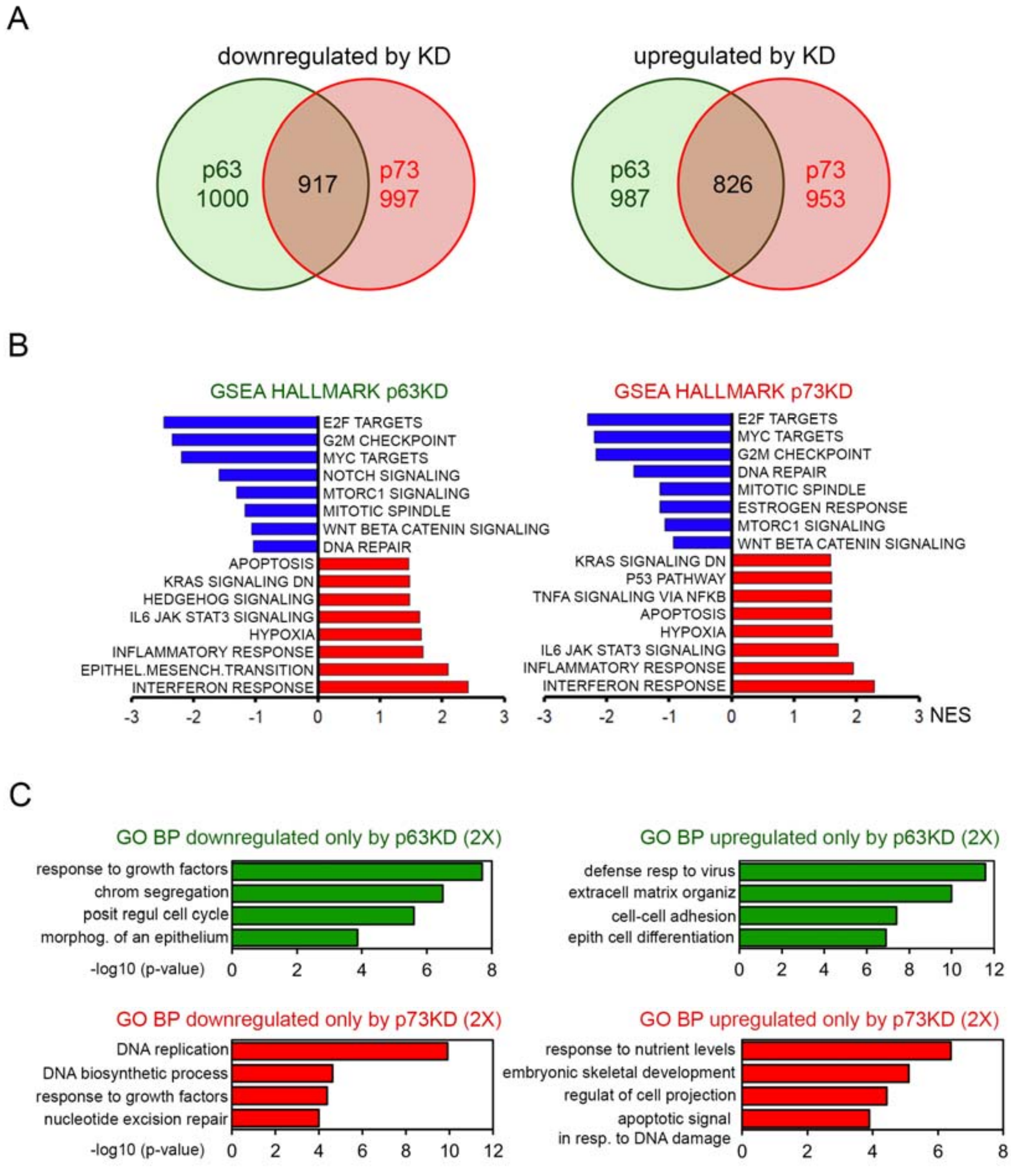


**Figure S3. Transcriptomic consequences of p63 and p73 depletion in SCC cells.** (A) Venn diagrams showing overlap of downregulated genes (left) and upregulated genes (right) upon p63 or p73 KD in SCC cells. (B) GSEA Hallmark analysis of differentially expressed genes following p63 (left) or p73 (right) depletion. Bars indicate significantly enriched gene sets among downregulated (blue) or upregulated (red) genes. X-axis: normalized enrichment score (NES). (C) GO Biological Process terms uniquely enriched upon p63 or p73 depletion. Terms were defined as specific based on ≥2-fold enrichment and FDR < 0.05 in one KD condition, with no significant enrichment in the other (FDR > 0.05).

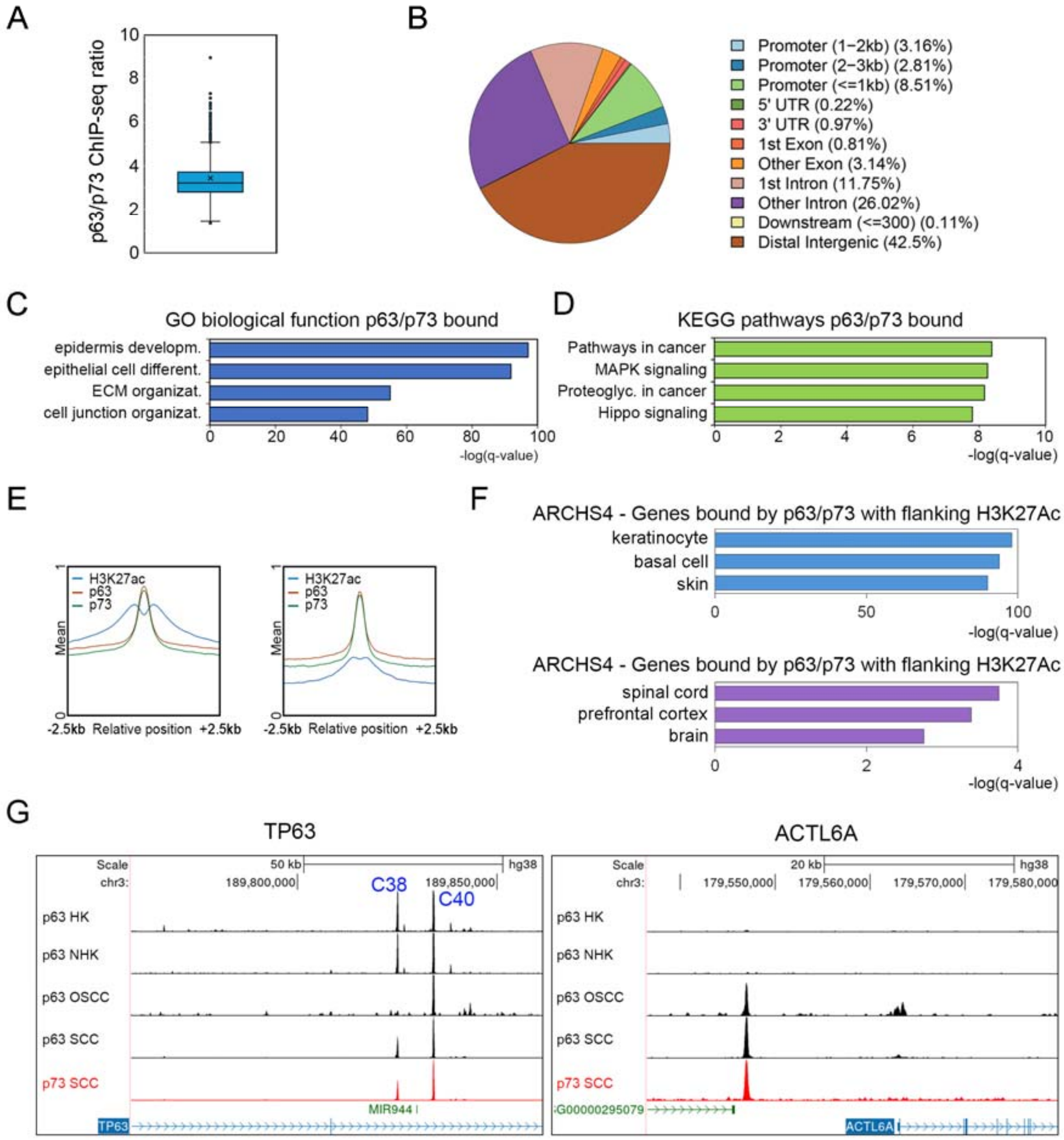


**Figure S4. Additional evidence for p63/p73 co-binding and context-dependent enhancer usage.** (A) Box plot showing the distribution of p63/p73 ChIP-seq signal ratios across co-occupied genomic regions. (B) Genomic distribution of p63/p73 co-bound ChIP-seq peaks assessed using ChIPseeker (105). (C) GO Biological Process enrichment analysis of genes associated with p63/p73 co-bound regions. (D) KEGG pathway analysis of p63/p73-bound genes. (E) Average ChIP-seq signal profiles (±2.5 kb) for p63, p73, and H3K27ac centered on p63/p73 co-bound peaks in SCC cells, indicating active regulatory elements. (F) ARCHS4-based tissue enrichment analysis (106) of genes linked to p63/p73 co-bound genomic regions flanked by H3K27Ac (top) or lacking flanking H3K27Ac (bottom).Bars represent −log(q-value). (G) Genome browser tracks illustrating representative examples of context-specific p63/p73 occupancy. Left: p63 locus showing p63 binding at conserved enhancers C38 and C40 in epidermal keratinocytes and SCC, with absence of C38 occupancy in OSCC. Right: *ACTL6A* locus showing strong p63/p73 binding in SCC and OSCC but not in keratinocytes.

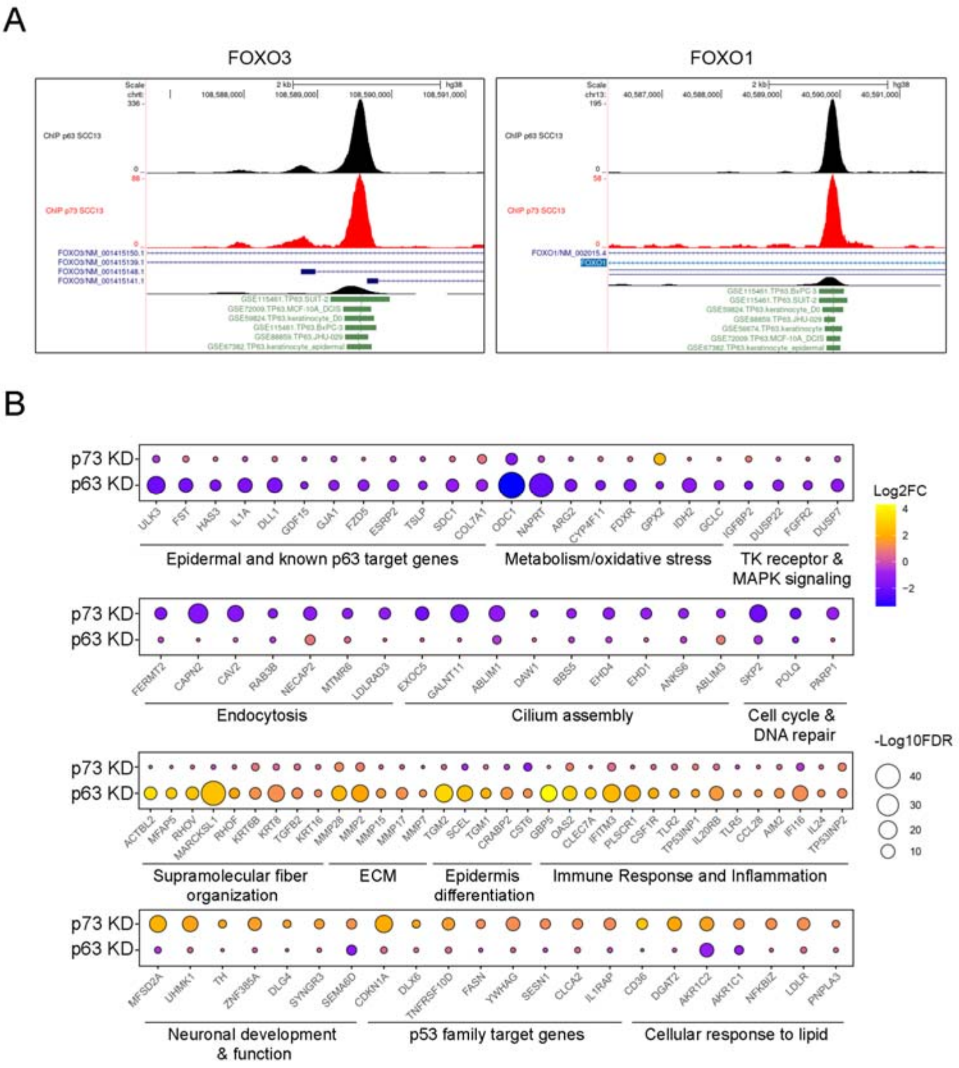


**Figure S5**. **Differential regulation of FOXO factors and pathway-specific target genes by p63 and p73.** (A) Genome browser tracks showing p63 (black) and p73 (red) ChIP-seq signal at the FOXO3 (left) and FOXO1 (right) loci in SCC cells. Intronic binding regions are co-occupied by both factors, consistent with direct regulation of FOXO genes across multiple epithelial contexts. including HNSCC (JHU-029), and pancreatic cancer cells (SUIT-2, BxPC-3)—based on ReMap datasets (107). (B) Balloon plot showing representative direct target genes differentially regulated by p63 or p73 KD across major functional categories. Balloon size represents −log10(FDR); color indicates log2 fold change.

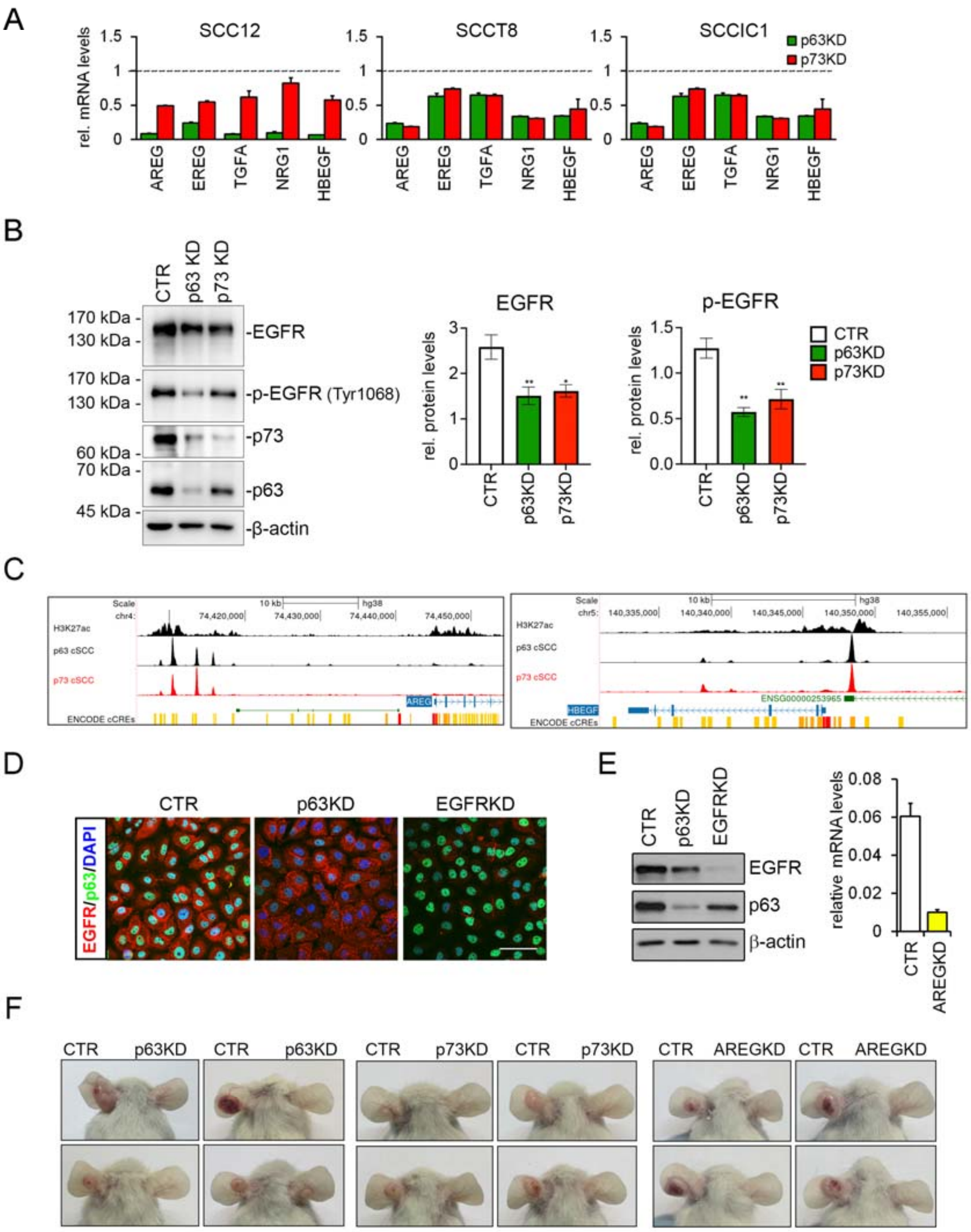


**Figure S6. Regulation of EGFR ligands and pathway activation by p63 and p73.** (A) RT-qPCR analysis of EGFR ligands mRNA levels in SCC cells after transfection with p63- or p73-specific siRNAs. Data are shown as mean ± SD relative to controls. (B) Immunoblot analysis (left) for the indicated proteins performed in SCC cells upon transfection of p63- or p73-specific siRNAs. Quantification (right) was performed using Image Lab (Bio-Rad). Data represent mean ± SD (n=3). Two-tailed Student's t-test: *P < 0.05; **P < 0.01. (C) Genome browser view of the H3K27ac-, p63-

and p73- ChIP-seq signals in the AREG (left) and HBEGF (right) gene loci. ENCODE cCRE tracks are shown. (D) Immunofluorescence analysis using antibodies specific for EGFR (in red) and p63 (in green) in SCC cells transfect with indicated siRNA or negative control. Nuclei were stained with 4,6-diamidino-2-phenylindole (DAPI; blue). Scale bar=50 μm. (E) Left: Immunoblot analysis of EGFR and p63 protein levels following indicated siRNA transfection. Right: RT-qPCR analysis of AREG mRNA levels under the same conditions. (F) Tumorigenic capacity of SCC cells assessed by ear injection in immunodeficient mice.
